# A pipeline to characterize local cortical folds by mapping them to human-interpretable shapes

**DOI:** 10.1101/2020.11.25.388785

**Authors:** Arnab Roy, Tyler McMillen, Donielle L Beiler, William Snyder, Marisa Patti, Vanessa Troiani

## Abstract

**Background:** Variations in regional cortical folds across individuals have been examined using computationally-derived morphological measures, or by manual characterization procedures that map distinct variants of a regional fold to a set of human-interpretable shapes. Although manual mapping approaches have proven useful for identifying morphological differences of clinical relevance, such procedures are subjective and not amenable to scaling.

**New Method:** We propose a 3-step pipeline to develop computational models of manual mapping. The steps are: represent regional folds as feature vectors, manually map each feature vector to a shape-variant that the underlying fold represents, and train classifiers to learn the mapping.

**Results:** For demonstration, we chose a 2D-problem of detecting within slice discontinuity of medial and lateral sulci of orbitofrontal cortex (OFC); the discontinuity may be visualized as a broken H-shaped pattern, and is fundamental to OFC-type-characterization. The classifiers predicted discontinuities with 86-95% test-accuracy.

**Comparison with Existing Methods:** There is no existing pipeline that automates a manual *characterization* process. For the current demonstration problem, we conduct multiple analyses using existing softwares to explain our design decisions, and present guidelines for using the pipeline to examine other regional folds using conventional or non-conventional morphometric measures.

**Conclusion:** We show that this pipeline can be useful for determining axial-slice discontinuity of sulci in the OFC and can learn structural-features that human-raters may rely on during manual-characterization.The pipeline can be used for examining other regional folds and may facilitate discovery of various statistically-reliable 2D or 3D human-interpretable shapes that are embedded throughout the brain.

## 1. Introduction

The human cerebral cortex is a sheet of neurons which varies in thickness (Economo, 1929; Fischl & Dale, 2000; Mai & Paxinos, 2012) and at a macroscopic level forms a convoluted structure of folding patterns of gyri and sulci (Borrell, 2018). A gyrus represents the convex portion of a cortical fold, whereas a sulcus represents the concave portion of the fold with white matter underneath (See Figure 1A) (Dahnke & Gaser, 2018; Yang & Kruggel, 2008). Although there is a general similarity across healthy individuals in the manner the cortical sheet folds (Duvernoy, 1992; Ono, Stefan, & Abernathey, 1990; Rhoton, 2007; ten Donkelaar, Tzourio-Mazoyer, & Mai, 2018; Wang, Necus, Kaiser, & Mota, 2016), subtle differences in folding patterns in specific brain regions are predictive of human behavioral traits (Amunts et al., 1997; Sun et al., 2012). Further, by examining regional variations in cortical folding patterns, studies have been able to differentiate patients with psychiatric disorders from controls (Bartholomeusz et al., 2013; Chakirova et al., 2010; Im, Raschle, Smith, Grant, & Gaab, 2016; Isomura et al., 2017; Lavoie et al., 2014; Li, Wang, Li, Li, & Li, 2015; Nakamura, Nestor, & Shenton, 2020; Patti & Troiani, 2018; Takayanagi et al., 2010). The above findings thus provide evidence that examining variations in sulcal folding patterns in a localized manner can have a broad range of clinical and neuroscience applications.

**Figure 1:**
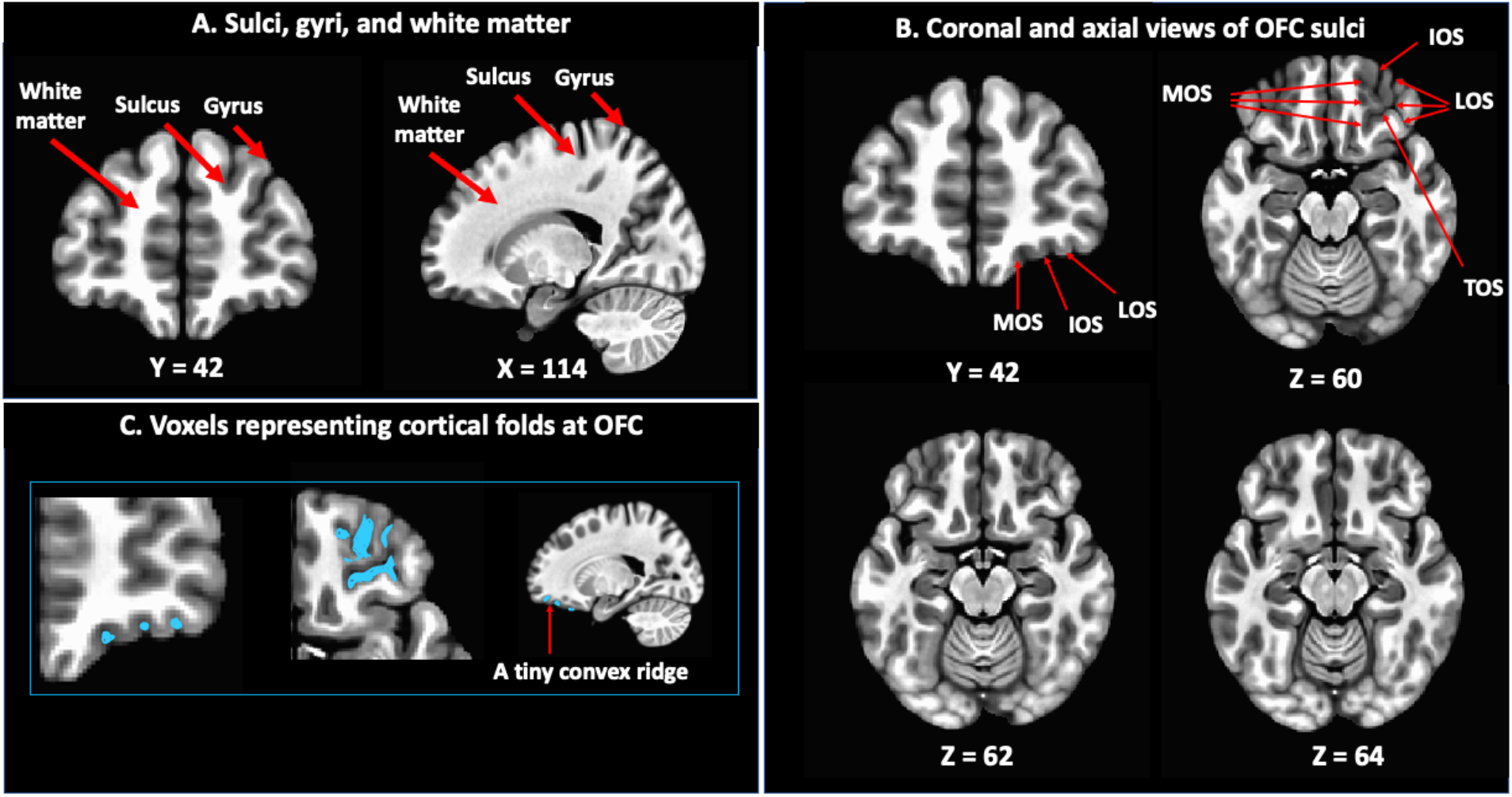
Gyrus, sulcus, and fragments of a cortical fold. **(A)** The figure illustrates an example of sulcus and gyrus using an averaged T1 MNI brain (*SSwarper base volumes*, 2020). **(B)** The figure shows how the cortical fold representing medial (MOS), intermediate (IOS), lateral (LOS), and transverse (TOS) orbital sulci vary across axial slices of an averaged T1 MNI brain. **(C)** The figure illustrates fragments of the cortical fold at OFC constituting MOS, IOS, LOS, and TOS.

### 1.1. Manual characterization of local cortical folds: advantages and challenges

Previous work has generally identified cortical sulci using anatomical atlases (Jaume, Macq, & Warfield, 2002; Sandor & Leahy, 1997; Thompson et al., 1997; Thompson & Toga, 1996) or supervised learning based procedures (Behnke et al., 2003; Goualher et al., 1999; Lohmann & Von Cramon, 2000, 1998; Rivière et al., 2002) and computationally examined their morphology using local curvature, sulcal depth, surface area, and/or number of gyral hinges (Li et al., 2009; Shimony et al., 2016; Li et al., 2017; Zhang et al., 2018). Alternately, investigators have manually examined and documented characteristic local cortical folds in healthy controls (Yousry et al., 1997; Chiavaras & Petrides, 2000; Mellerio et al., 2016; Zhang, Harris, Split, Troiani, & Olson, 2016), in patients (Bartholomeusz et al., 2013; Isomura et al., 2017; Lavoie et al., 2014; Nakamura et al., 2020; Patti & Troiani, 2018; Takayanagi et al., 2010; Wang et al., 2020; Whittle et al., 2014; Zhang et al., 2016), or in brains obtained at autopsy (White et al., 1997; Zhang et al., 2010, 2011). In much of this work, characterization involved a manual documentation strategy, whereby, local cortical folds were mapped to a set of human-interpretable shapes and the mappings were summarized using pictorial illustrations (Yousry et al., 1997; Chiavaras & Petrides, 2000; Mangin et al., 2019; Mellerio et al., 2016; Weiner et al., 2014). For example, Weiner and colleagues (2014) identified a reliable ω-shaped pattern in the mid-fusiform sulcus, when seen from a coronal view, and further categorized the sulcus into four subtypes each representing a variant of the ω-shape. Similarly, Chiavaras and Petrides (2000) documented multiple sub-types of sulcal patterns within the orbitofrontal cortex (OFC) each representing a specific variant of the letter “H”. In all, the above mapping approach simplifies visualization of how patterns of regional cortical folds may vary and facilitates manual characterization of such folds.

Despite the continued use of these manual sulcal characterization procedures to reveal interesting neurobiological insights with clinical relevance (Bartholomeusz et al., 2013; Isomura et al., 2017; Lavoie et al., 2014; Nakamura et al., 2020; Patti & Troiani, 2018; Takayanagi et al., 2010; Wang et al., 2020; Whittle et al., 2014; Zhang et al., 2016), there are a number of problems that limit their widespread application. The procedures involve a degree of subjectivity, as illustrated by interrater reliabilities scores between 0.71 to 0.89 (Bartholomeusz et al., 2013; Chakirova et al., 2010; Isomura et al., 2017; Lavoie et al., 2014; Li et al., 2019; Nakamura et al., 2020; Patti & Troiani, 2018; Takayanagi et al., 2010; Wang et al., 2020; Whittle et al., 2014; Zhang et al., 2016). The manual characterization itself is also a slow and arduous process, and is not suited for examining big datasets. Also, while developing manual guidelines to map regional cortical folds to a predefined set of human interpretable shapes, it is very difficult to concisely and accurately explain the variability in this mapping, which thereby impedes the discovery of other embedded shapes in regional cortical folds across the brain. One way to overcome the above challenges is to develop a computational framework that uses example cases characterized by human raters that represent the mapping between variations of a regional cortical fold and a set of human interpretable shapes. This data can then be used for training a classification system that can then be used to predict the mapping for new cases. Such a framework would eliminate rater subjectivity, enable processing larger datasets, and will serve as a tool for investigators to reliably test the embedded shapes in other cortical areas. In this work, we introduce such a framework, and demonstrate the approach, using Chiavaras and Petrides’ (2000) broken H-pattern detection problem, which we discuss below.

### 1.2. The H-pattern detection problem: computational challenges and utility

The shape of the cortical fold within the OFC generally represents an H-shaped pattern consisting of four sulci: medial (MOS), transverse (TOS), lateral (LOS), and intermediate (IOS) orbital sulci (See Figure 1B). As shown in Figure 1B, MOS and LOS represents the vertical arms of the H-shaped pattern, and the TOS represents the horizontal line connecting these vertical arms. Chiavaras and Petrides (2000) first described this “H” pattern in structural MRI images of 50 healthy human subjects along with descriptions of distinct variations of the H-pattern. That is, the “H” pattern variations were differentiated based on whether the MOS or LOS arms were discontinuous (broken) within an OFC axial slice. An example of a discontinuous MOS and LOS is shown in Figure 1C, where the blue colored segments representing the cortical folds at the MOS and LOS are spatially disconnected. Based on identifying this discontinuity, we and others (Bartholomeusz et al., 2013; Chakirova et al., 2010; Isomura et al., 2017; Lavoie et al., 2014; Li et al., 2019; Nakamura et al., 2020; Patti & Troiani, 2018; Takayanagi et al., 2010; Troiani, Patti, & Adamson, 2019; Wang et al., 2020; Whittle et al., 2014; Zhang et al., 2016) have manually characterized the “H” sulcus into 4 pattern types: Type I is a discontinuous MOS and continuous LOS, Type II is a continuous LOS and MOS, Type III is a discontinuous MOS and LOS, and Type IV is a continuous MOS and discontinuous LOS, and found the distribution of the above pattern types to be atypical in groups of subjects with neuropsychiatric illness. Characterizing the discontinuity of MOS and LOS within an axial slice depends on carefully examining the sizes and the spatial organization of the individual MOS or LOS fragments relative to each other and also relative to the TOS. The characterization remains challenging, given the large degree of variability in the H-sulcus across subjects (see Figure 2). Thus instead of addressing the issue of type characterization, we directly address the fundamental issue of how to automate the process of identifying MOS and/or LOS discontinuity (broken-ness) within an axial slice, which represents the broken vertical arm of an H-shaped pattern.

**Figure 2:**
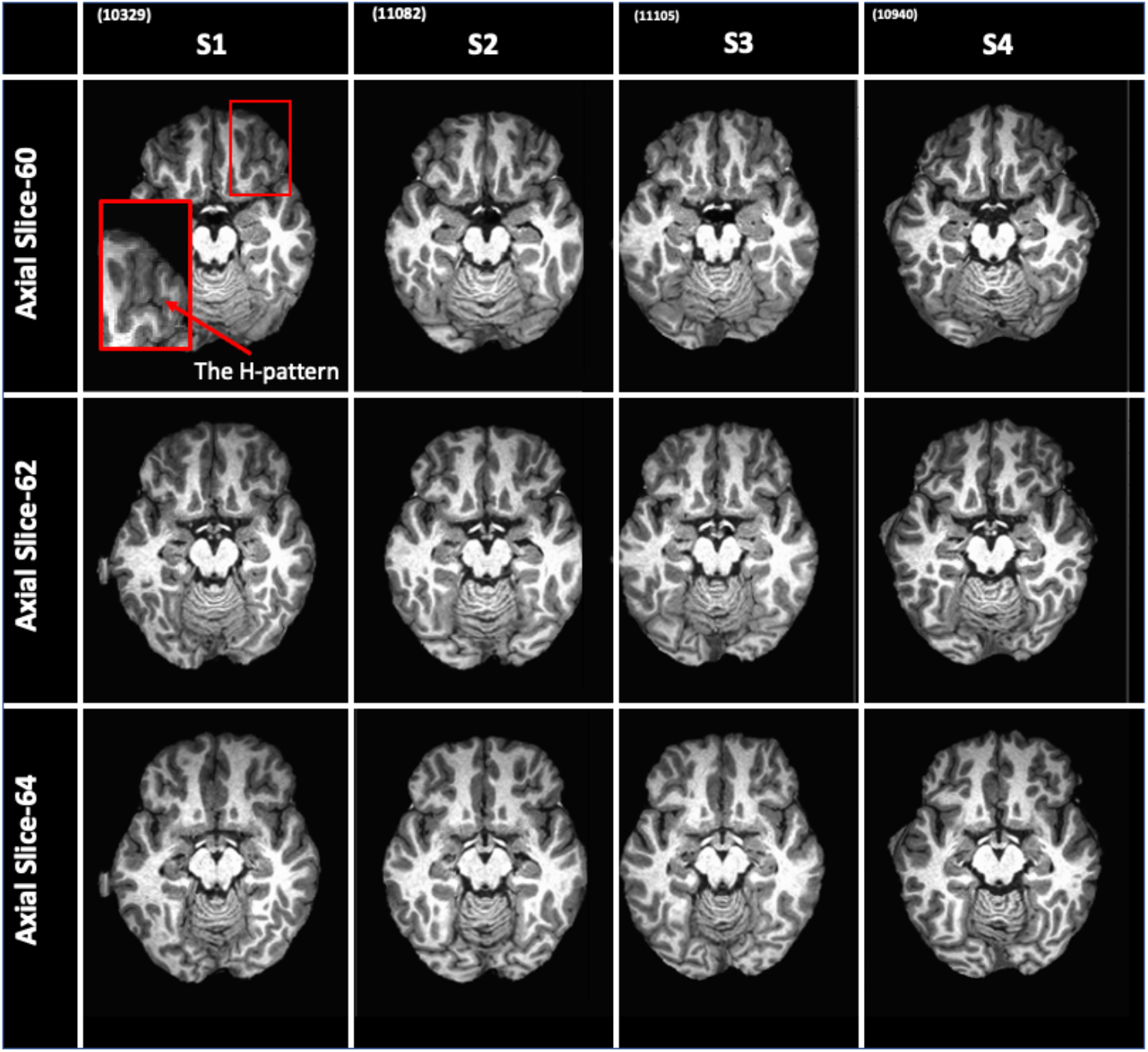
The H-pattern. The figure illustrates how the sulcus at OFC resembles a H-shaped pattern when seen from an axial view. That is, for a given OFC axial slice, for the right hemisphere, the MOS represents the left vertical line of the letter “H”, the LOS represents the right vertical line, and the TOS represents the horizontal line connecting MOS and LOS. Similarly, for the left hemisphere, the MOS represents the right vertical arm of the letter “H” and LOS represents the left vertical arm. The pattern varies across axial slices of a subject, and also there is inter-subject variance.

### 1.3. An overview of our framework

Broadly, the steps in our computational framework involve encoding regional cortical folds as feature vectors, assigning a numeric label to each feature vector depending on the human-interpretable shape the cortical fold it encodes represents, and then training a classifier using N-fold crossvalidation to learn this mapping. Although, any supervised-classification technology may be integrated within our framework, here we used a fixed architecture neural network classifier. We also chose to create feature vectors within 2D axial slices in order to directly mimic the steps followed by commonly used manual characterization procedures within the H-sulcus (Bartholomeusz et al., 2013; Chakirova et al., 2010; Isomura et al., 2017; Lavoie et al., 2014; Li et al., 2019; Nakamura et al., 2020; Patti & Troiani, 2018; Takayanagi et al., 2010; Troiani et al., 2019; Wang et al., 2020; Whittle et al., 2014; Zhang et al., 2016). However, alternate strategies may be used for developing feature vectors using 3D tissue segmentation and mesh-based approaches, and their elements may represent conventional regional morphometric measures (Auzias et al., 2014; Borne, 2019; Duan et al., 2019; Jin et al., 2020; J. F. Mangin et al., 2015) and/or measures capturing interregional morphometric coherence (Corps & Rekik, 2019; Nebli & Rekik, 2019; Zheng et al., 2018). In all, the framework may be flexibly used for characterizing cortical folds within other brain regions.

## 2. Methods

### 2.1. Participants

The structural images used in this study were obtained from the Openfmri database (Bilder et al., 2018). Here, we used 52 control subjects from this dataset whose OFC were examined in a recent publication using manual tracing (Patti & Troiani, 2018). All subjects gave written and informed consent to the Institutional Review Board at UCLA.

### 2.2. Training classifiers to learn MOS and LOS discontinuity within OFC axial slices

The current implementation of the framework, which is illustrated in Figure 3, consisted of the following processing stages: (A) quality control and normalization of T1 images, (B) creating MOS, TOS, LOS, and IOS probability maps using an independent dataset so that voxels representing OFC sulci may be identified, (C) encoding the spatial organization of pairs of MOS and LOS sulcal-fragments within axial slices as feature vectors, (D) manually assigning labels to the feature vectors based on traditionally used guidelines and (E) training neural network classifiers to learn the mapping. We discuss each step in detail, below.

**Figure 3:**
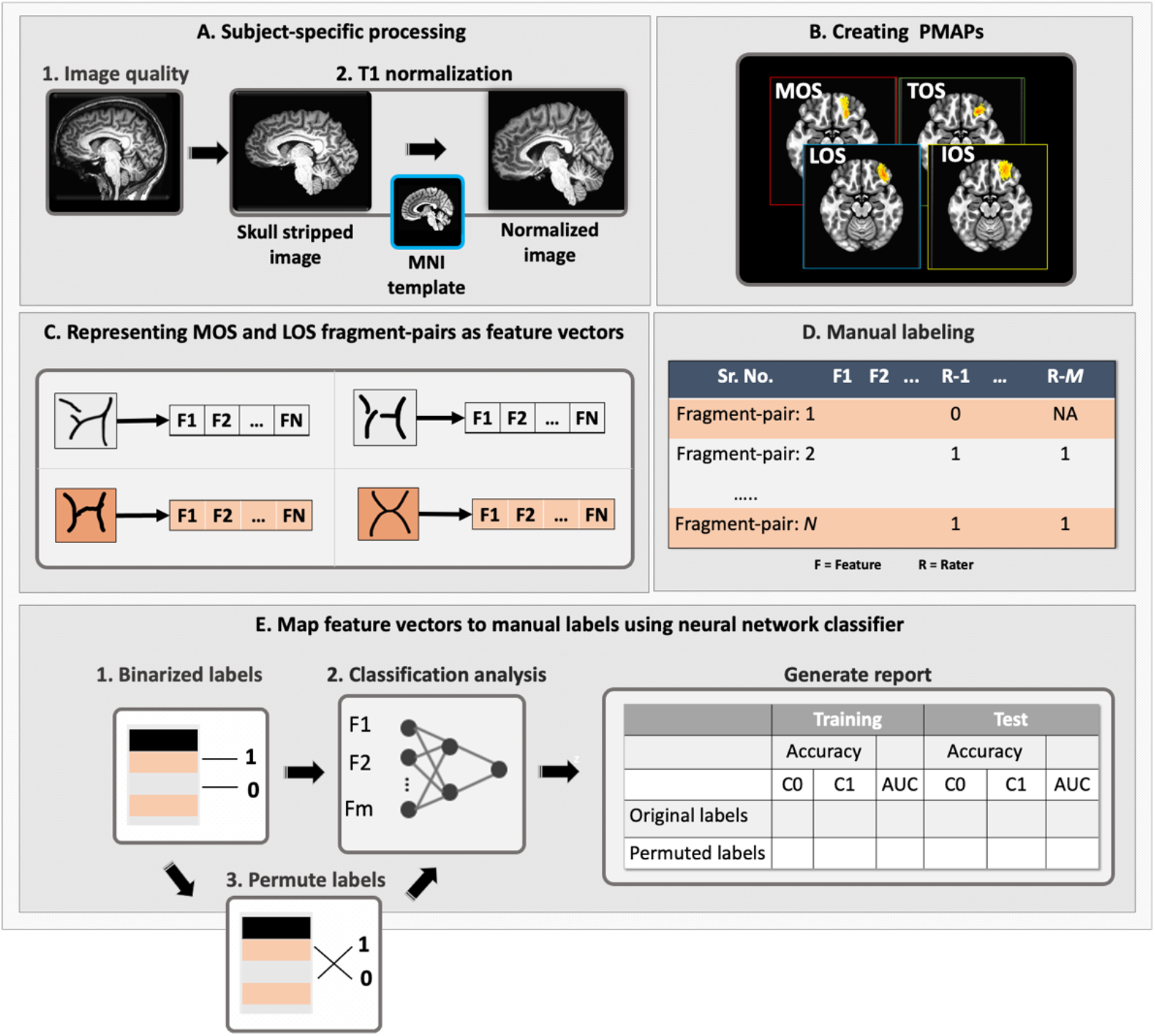
The analysis pipeline. The figure illustrates the processing stages of our computational framework for characterizing MOS and LOS within axial slice discontinuity. **(A)** In the first stage, we normalize an individual’s T1-weighted image to MNI space and exclude noisy images. **(B)** In the second stage, MOS, TOS, LOS, and IOS, PMAPs are created using an independent dataset (Troiani et al., 2019) using an established permutation based procedure (Weiner et al., 2017) which are then used for identifying the voxels corresponding to the above sulci in each subject’s OFC slice. **(C)** In the third stage, pairs of MOS and LOS fragments are encoded as feature vectors (See Figure 6 for more details). **(D)** In the fourth stage, the feature vectors are manually labeled by multiple raters as 0 or 1, depending on MOS or LOS discontinuity. **(E)** In the final stage, using neural network based classification analysis, the mapping between the feature vectors and the labels is established.

#### 2.2.1. Subject-specific processing

We used the MRI Quality Control (MRIQC) toolbox (Esteban et al., 2017) to identify and discard subjects whose T1 image was contaminated by motion and intensity nonuniformity artifacts. We discarded subjects (N=3) with a coefficient of joint variation above 0.5, which indicates a greater contamination of artifacts (Esteban et al., 2017; Zin, Zheng, Chee, & Zagorodnov, 2009; See Supplemental Table S1). The cases (N=49) that passed MRIQC were normalized to MNI space (See Figure 3A) using AFNI software (Cox, 1996). The normalization process involved removing shading artifacts, skull-stripping, affine transformation to MNI-space, and non-linear warping to an AFNI template in MNI space; see Supplemental Section I.a for additional details. The normalized images were visually inspected for distortions (e.g., over smoothing, stretching) within the OFC and N=2 cases were discarded from further analysis due to distortions.

The results of the existing literature suggest that cortical surface area and cortical volume are positively correlated, at both whole-brain and regional levels (Toro et al., 2008; Winkler et al., 2010), and as the size of cortex increases, so does the degree of sulcal convolution (Im et al., 2008). To ensure that the results of our classification analyses are not influenced by subjects with unusually high or low OFC gray matter volume, we conducted a box plot analysis using whole-brain and local OFC volumetric measures, and found no outlier cases. In Supplemental Section I.b, we have discussed the above computational steps in detail, and in Supplemental section II.a and Supplemental Table S2 we have reported all subject’s volumetric data.

#### 2.2.2. Creating MOS, TOS, LOS, and IOS probability maps using an independent dataset

We generated probability maps (PMAPs) of left and right MOS, TOS, LOS, and IOS (See Figure 3B), by running an established permutation-based averaging procedure (Weiner et al., 2017) using an independent dataset of manual OFC tracings from 47 subjects (Troiani et al., 2019). In the manual tracing file of each subject, the voxels belonging to each OFC sulcus were represented by a unique numeric ID. We first identified a sulcus using its corresponding ID, and then binarized all the voxels that were part of the sulcus. We then averaged each subject’s binarized sulcal tracing to generate an initial PMAP of each sulcus. The value within each voxel thus represented the proportion of subjects that overlapped at that voxel for a given sulcus. We then used a leave-one-out cross-validation procedure, described elsewhere (Weiner et al., 2017), to determine an optimal probability threshold that produced the most predictive sulcal map for each sulcus. That is, at the optimal threshold, the overlap between the training PMAPs and the test set PMAP (the leftout case), measured as dice coefficient, was maximum. All voxels in the initial averaged PMAP of a sulcus with values lower than the corresponding optimal threshold were set to 0 and the resulting image represented the final averaged PMAP for the sulcus. The final PMAPs were transformed from FSL’s MNI-space (space in which manual tracings were available) to AFNI’s MNI space using an affine transformation (i.e., 3dAllineate function). We chose affine over non-linear transformation to minimize the regional distortion of probability values encoded within the PMAPs. The results of the PMAP analysis are summarized in Figure 4 and Figure 5. As can be seen in the figures, the dice coefficient values ranged from 0.14 to 0.17, thus suggesting some level of intersubject variability in the manner the cortex folds within the OFC.

**Figure 4:**
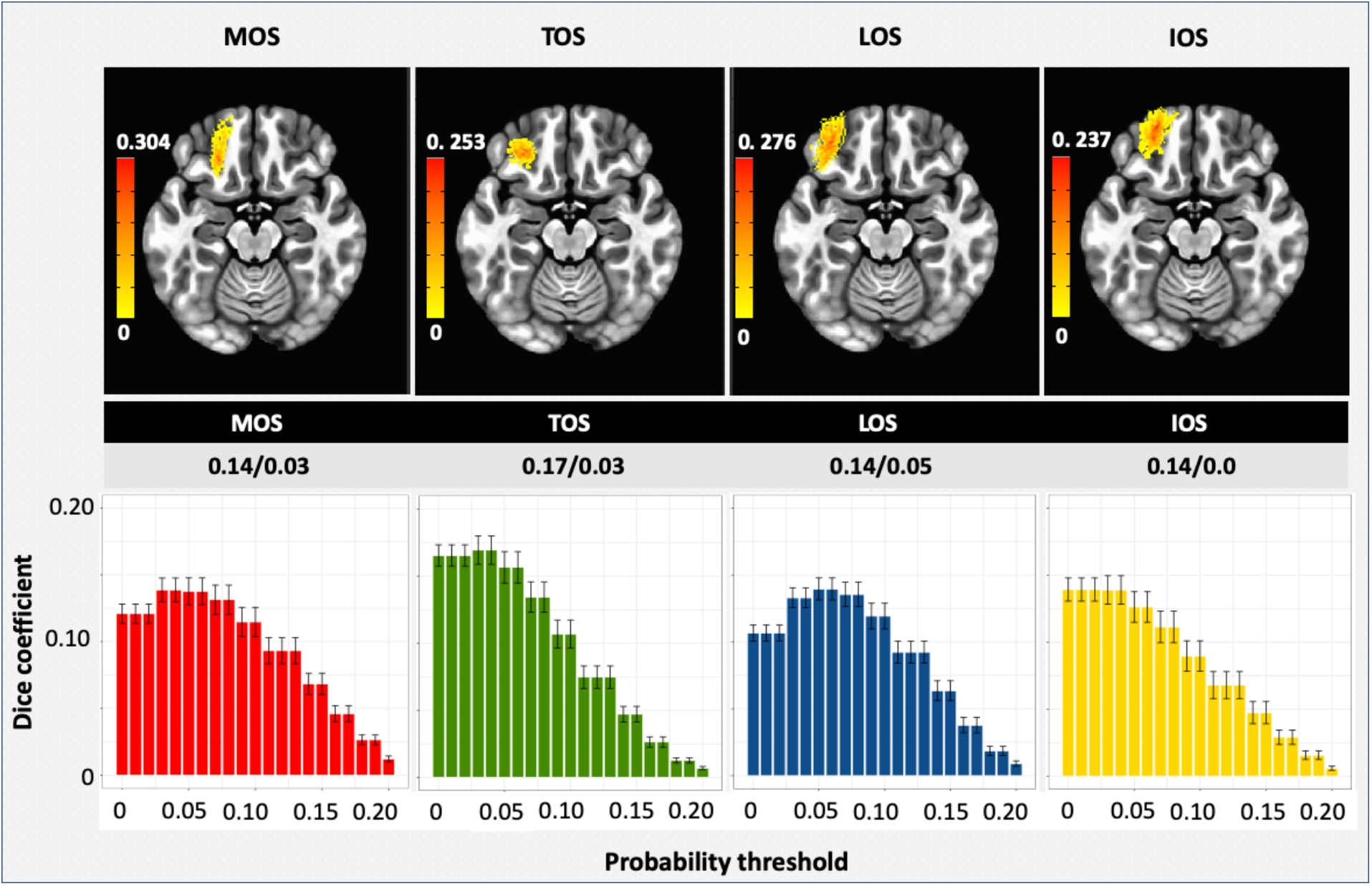
PMAPs for left OFC sulci. In the top row, we illustrate the left OFC PMAPs that we created using an independent dataset consisting of manual OFC sulci tracings (Troiani et al., 2019). In the bottom row, moving from left to right, we illustrate the dice coefficient versus probability threshold graphs for MOS (red), TOS (green), LOS (blue), and IOS (yellow). The graphs were generated by the leave-one-out type permutation procedure for establishing an optimal probability threshold for developing the final PMAPs. For MOS, the optimal probability threshold was 0.03 with a dice coefficient of 0.14, for TOS the optimal probability threshold was 0.03 with a dice coefficient of 0.17, for LOS the optimal probability threshold was 0.05 with a dice coefficient of 0.14, and for IOS the optimal probability threshold was 0.00 with a dice coefficient of 0.14.

**Figure 5:**
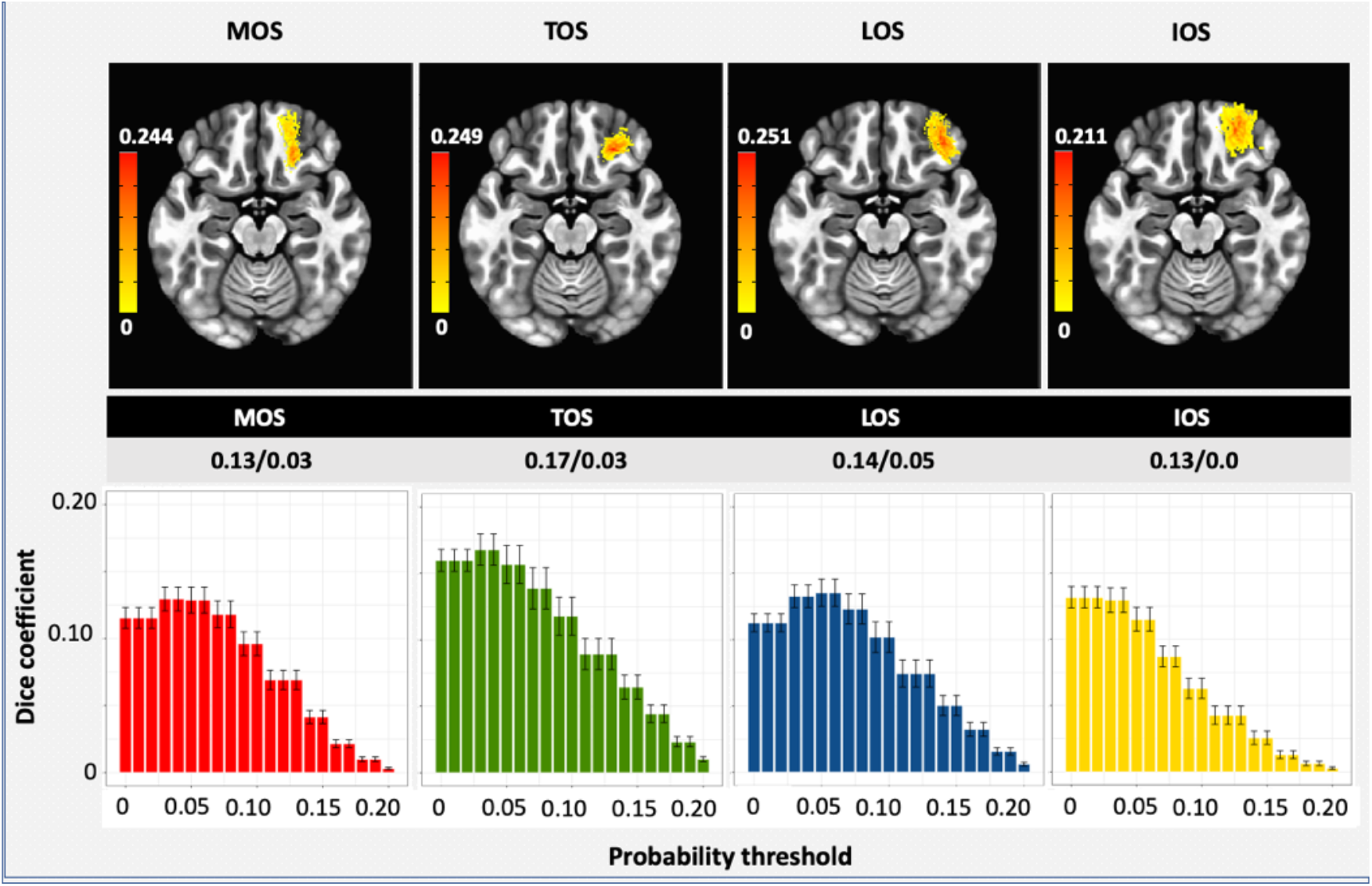
PMAPs for right OFC sulci. In the top row, we illustrate the left OFC PMAPs that we created using an independent dataset consisting of manual OFC sulci tracings (Troiani et al., 2019). In the bottom row, moving from left to right, we illustrate the dice coefficient versus probability threshold graphs for MOS (red), TOS (green), LOS (blue), and IOS (yellow). The graphs were generated by the leave-one-out type permutation procedure for establishing an optimal probability threshold for developing the final PMAPs. For MOS, the optimal probability threshold was 0.03 with a dice coefficient of 0.13, for TOS the optimal probability threshold was 0.03 with a dice coefficient of 0.17, for LOS the optimal probability threshold was 0.05 with a dice coefficient of 0.14, and for IOS the optimal probability threshold was 0.00 with a dice coefficient of 0.13.

#### 2.2.3. Encoding the spatial organization of pairs of MOS and LOS fragments as feature vectors

We computationally mimicked the established and most ubiquitously used manual H-sulcus tracing procedure (Bartholomeusz et al., 2013; Chakirova et al., 2010; Chiavaras & Petrides, 2000; Isomura et al., 2017; Lavoie et al., 2014; Li et al., 2019; Nakamura et al., 2020; Patti & Troiani, 2018; Takayanagi et al., 2010; Troiani et al., 2019; Whittle et al., 2014; Zhang et al., 2016) to create feature vectors to represent MOS or LOS discontinuity within OFC axial slices of a subject (Figure 3C). Broadly, there were two steps: fragments of cortical folds representing MOS, TOS, and LOS were identified and then a feature vector was created for each pair of MOS (or LOS) fragments based on their size and spatial organization. We discuss these steps in detail below:

##### A. Identifying fragments of OFC cortical fold

Slice-specific processing was run using AFNI normalized T1 images of the subjects who passed image quality analyses. In previous work by our lab (Snyder, Patti, & Troiani, 2019), we found that individual OFC sulci were not always captured using existing software for automated sulcal tracing, such as BrainVisa. Importantly, the commonly available segmentation tools for segmenting gray matter, white matter, and CSF are not necessarily useful for identifying this sulcal shape, as the manual tracing procedure tends to include both CSF and gray matter voxels. For this reason, we developed a simple 2D procedure to identify voxels that are part of the OFC cortical fold representing MOS, TOS, LOS, and IOS using an adaptive histogram-based approach; similar histogram-based procedures are common in medical image analysis (Choi et al., 2016; Fujima et al., 2019; Li et al., 2019).

The procedure first extracts the axial slices 58 to 68 from each subject’s T1 normalized image using a custom-made program which utilizes R’s fMRI package (Tabelow & Polzehl, 2011). As can be seen in Figure 6A, the voxels that represent the sulci have much lower image intensity than surrounding voxels, and hence, they may be easily identified computationally by setting an intensity threshold. To allow such thresholds to vary across axial slices and subjects, the procedure first creates a histogram of voxel intensity values for each axial slice, and the image intensity at the 90^th^ percentile bin is identified. The threshold is then defined as 0.65 times the 90^th^ percentile image intensity value; we set this multiplication factor to 0.65 based on preliminary visual tests. All voxels with a value less than the threshold and within a rectangular box surrounding the OFC, defined using anatomical guidance, are then identified as voxels representing cortical folds in OFC (See Figure 6B, cyan cluster). As the regions near the brain-boundary have near 0 values, the histogram-based procedure also picks-up these voxels (See Figure 6: boundary identified in pink). We discarded such boundary voxels at OFC using a mask, which we created as follows.

**Figure 6:**
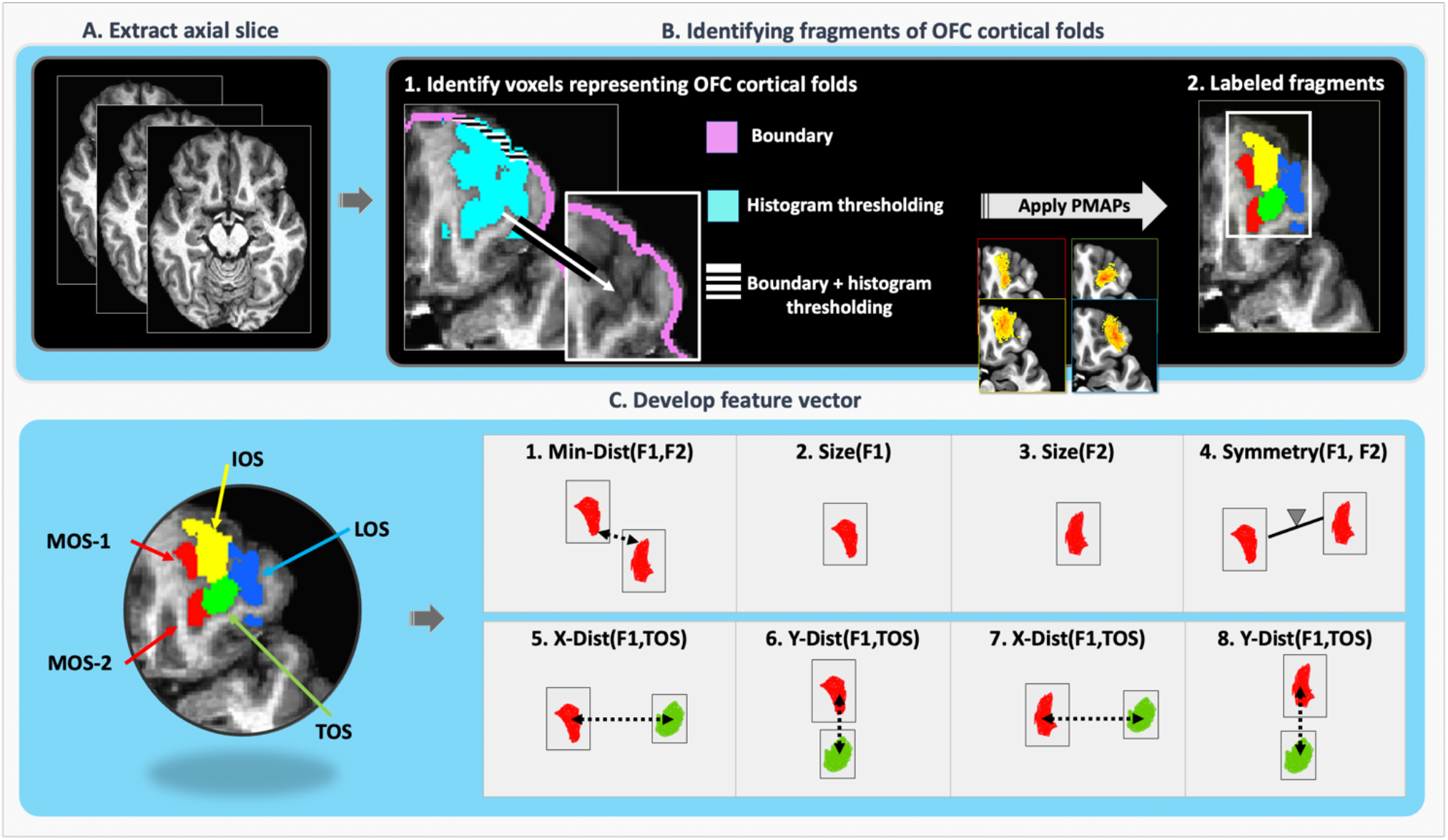
The feature vector. The figure illustrates the steps we followed for encoding the spatial organization of a pair of MOS or LOS fragments within an axial slice as a feature vector. Broadly, there were three steps: **(A)** extracting an OFC axial slice, **(B)** identifying voxels that represented MOS, TOS, LOS, and IOS in the axial slice using an adaptive histogram-based procedure, assigning labels to the voxels using PMAPs and grouping them into spatially disconnected fragments, and **(C)** encoding the spatial organization of a pair of MOS or LOS fragment, as an 8-element feature vector.

We first converted AFNI’s MNI template to a binary image. For each axial slice of this mask, we then identified the boundary voxels by first choosing a y-axis value (the y-axis represented the posterior to anterior coordinate), and then scanning the image by starting from the leftmost extreme, which always represented out-of-brain region, and moving towards the right, and stopping when the first voxel with a value of 1 was detected representing a brain-boundary voxel. The above scanning procedure was run for all possible y-axis values. In a similar manner, the procedure was run from right to left, and from anterior to posterior directions. The identified boundary voxels were discarded from the original mask, and the above procedure was run once again, yielding another layer of boundary voxels. The identified 2-layered boundary voxels are illustrated in Figure 6B: Step-1. The voxels identified by the histogram-based procedure that did not overlap with this 2-layered boundary, and were strictly within it, were saved for further analysis. The current strategy was suitable given the brain-boundary at OFC represents a convex shape, although alternate identification of brain-boundary voxels can certainly be used.

To label the surviving voxels as MOS, TOS, LOS, and IOS, we overlaid the corresponding axial slice of the 4 PMAPs (See Section 2.2.2) for each OFC axial slice for each subject, separately for each hemisphere. A voxel was labeled as 1 if the MOS PMAP’s value at the voxel was greater than the other PMAP values. Similarly, a voxel was labeled as 2 if the LOS PMAP’s value at the voxel was the highest, 3 if the TOS PMAP’s value at the voxel was highest, and 4 if the IOS PMAP’s value at the voxel was the highest. All voxels for which the PMAP value was 0 were discarded from further analyses. We then obtained fragments of cortical folds representing MOS, TOS, and LOS by first creating a separate lattice graph (each voxel at most shared edges with 4 spatial neighbors) for each sulcus, and then identifying the voxels that constituted the disjoint components of the graph (See Figure 6B: Step-2). We conducted an additional analysis to quantify the percentage of cerebrospinal fluid (CSF), gray matter (GM), and white matter (WM) within the sulcal-fragments identified using the above procedure. Our results suggested that on average ~36% (sd=13%) of voxels within these fragments were CSF, ~63% (sd =13%) were GM, and 0% (sd = 0%) were white matter. In all, the sulcal-fragments were primarily composed of CSF and gray matter, consistent with the voxel tissue types captured using manual tracing procedures.For some cases, mostly in the inferior slices, there were a few mislabeled fragments that existed slightly outside the OFC. We visually identified these fragments using anatomical guidance and excluded them from the classification analyses. Additionally, some voxels were mislabeled near regions representing MOS-TOS, TOS-LOS, MOS-IOS, and IOS-LOS boundaries. Such boundary voxel mislabeling might have introduced some noise in our estimation of individual elements of the feature vectors, which we discuss below. Also, for some rare cases, the mislabeling led to fragmentation of MOS or LOS near the TOS junction, when visually it seemed no such discontinuity existed. Based on independent judgement of multiple raters, we discarded fragment-pairs that should not have been fragmented. But mislabeling that existed near sulcal boundaries that merely would lead to mis-estimation of a fragment’s size, pair-distance, and or symmetry were still included, as we anticipated inclusion of some noise might be good for developing a robust classifier.

##### B. Developing sulcal-fragment based feature vectors

Based on the typical human characterization procedure used by our lab and others (Bartholomeusz et al., 2013; Chakirova et al., 2010; Isomura et al., 2017; Lavoie et al., 2014; Li et al., 2019; Nakamura et al., 2020; Patti & Troiani, 2018; Takayanagi et al., 2010; Troiani et al., 2019; Wang et al., 2020; Whittle et al., 2014; Zhang et al., 2016), discontinuity between a pair of MOS or LOS sulcal-fragments can be determined using several criteria. These criteria can be generalized as follows: both fragments must be reasonably large and close to each other, their sizes well-balanced, both fragments are reasonably close to the TOS and are well-aligned in the y-direction, and one fragment is reasonably posterior to the TOS, and the other is in a relatively anterior direction. For each pair of sulcal-fragments, the above factors were encoded as an 8-element feature vector. In Figure 6C, we illustrate the elements using a pair of MOS sulcal-fragments; the same strategy was used for encoding the spatial organization of a pair of LOS sulcal-fragments. As shown in Figure 6C, element-1 encoded the minimum Euclidean distance between the voxels of the two MOS fragments, elements 2 and 3 encoded the sizes of individual MOS sulcal-fragments in voxels, element-4 represented the size symmetry of the two MOS fragments using equation 1, elements 5 and 6 represented the distance of the first MOS fragment’s centroid from TOS centroid in x- and y-directions, respectively, and elements 7 and 8 represented the distance of the second MOS fragment’s centroid from TOS centroid in x- and y-directions, respectively; the centroid of each cluster was estimated as the mean x- and y-axis values of the cluster estimated using the constituent voxels. We will refer to a feature vector consisting of these eight features as, *FV*_*fragment*_.

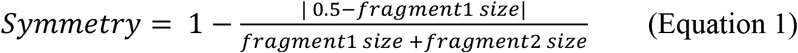

##### C. Developing spine based feature vectors

A common practice to manually characterize the H-sulcus is to use thin curved lines to represent the OFC sulci. Although, such a line representation may be computationally derived using existing pipelines like BrainVISA’s morphologist (Fischer et al., 2012), a published study by our lab (Snyder et al., 2019) found that when using the pipeline, several sections of the OFC sulci were underrepresented. We found the same effect when we ran BrainVISA’s Morphologist-2015 pipeline on the AFNI normalized T1 images of our subjects to derive their sulcal skeletons (See Table S3); the morphologist pipeline was able to generate a line representation only for about 55%-65% of the left/right LOS and MOS sulcal-fragments. We anticipate that this effect might be due to the fact that line representations in 3D mesh space are based on identifying a path across mesh-nodes that represent image minimas across OFC axial slices, and might therefore, under-represent image minimas within axial slices. Hence, we developed a custom-made procedure to establish a line representation of individual fragments of cortical folds representing MOS, TOS, and LOS within axial slice by directly using the data in voxel space (instead of mesh space), which we refer to as spines (See Figure 7 and Supplemental Section I.c); the approach generated spines for 99% of the sulcal-fragments (See Table S3). A supplemental analysis using sulcal-fragments for which both spines and BrainVisa line representation could be established, suggested that both approaches identified spatially similar set of voxels of low intensity that together represented the general structure of the fold (See Supplemental Sections I.d and II.b).

**Figure 7:**
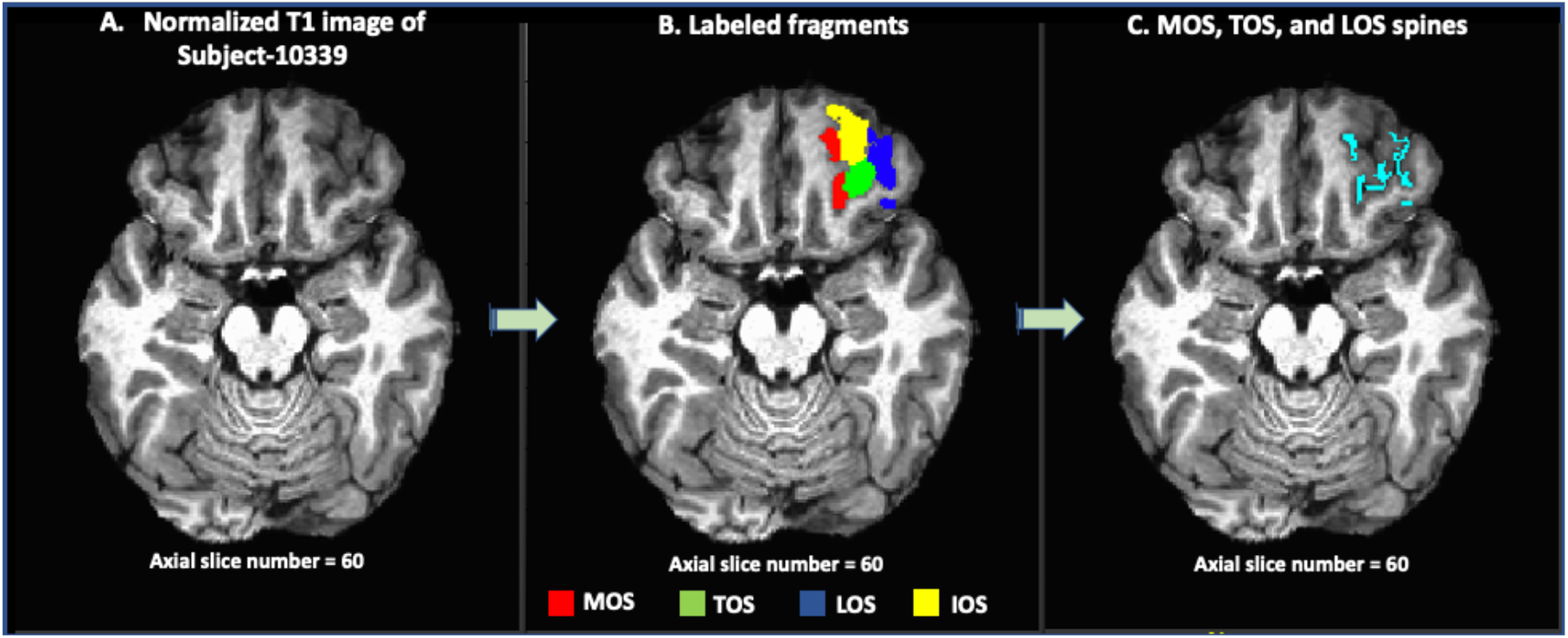
Labeled fragments and their spines. **(A)** The figure illustrates an axial slice of subject’s normalized T1-weighted image. **(B)** The MOS fragments are shown in red, IOS fragments in yellow, TOS fragments in green, and LOS fragments in blue. **(C)** The figure illustrates the spine of MOS, TOS, and LOS fragments identified using our custom-made spine detection technique.

Using the spines, we then developed a similar 8-element feature vector, *FV*_*spine*_, per fragment-pair using their spines. That is, element-1 of the feature vector represented the minimum distance between the spines of the two fragments, the element-2 and element-3 represented the sizes of the fragment’s spines, and element-4 represented the symmetry of their sizes. The final four elements were the same as *FV*_*fragment*_ features, as they represented the x and y-axis distances of the fragment’s centroids from TOS centroid. Although, we could have computed such centroids for spines, and then evaluated their x- and y-axis distances from TOS centroids, we relied on the fragment’s centroid instead because for some fragments the number of voxels in their spine were too few to reliably estimate a centroid. All *FV*_*fragment*_ and *FV*_*spine*_ feature vectors across subjects for each sulci, were saved in *FV*_*fragment*_ and *FV*_*spine*_ feature tables, repectively. The above steps yielded 8 feature tables (2 hemisphere x 2 sulci x 2 feature vector types) so that 8 classification analyses may be run to gauge the effetiveness of these features, and also to examine if feature vectors developed from spines were as effective as feature vectors developed using the fragments themselves.

#### 2.2.4. Mapping feature vectors to manual labels using neural network classifiers

The overall objective of the classifier was to learn to predict discontinuity of MOS or LOS within an OFC axial slice using size and distance-based features of fragment-pairs. The classifier utilized a 5-fold cross-validation type supervised learning framework. The procedure was run separately for left MOS, right MOS, left LOS, and right LOS. We discuss these steps below for right MOS; the same steps were applied for processing left MOS, left LOS, and right LOS.

##### Labeling of MOS and LOS fragment-pairs

For each subject, for each axial slice, the right MOS fragments were plotted and saved as an image file. Four raters (AR, TM, DB, and WS) were trained to identify discontinuous MOS by examining pairs of MOS fragments using a separate dataset of manual tracings. The raters then independently inspected each pair of fragments (and also their spines) and labeled the pairs as 0, 1, or NA. For a given fragment-pair, a rater labeled the fragment-pair as 1 if the MOS within the slice was discontinuous (while excluding all other fragments). If the discontinuity could not be determined, the fragment-pair was labeled as 0 (See Figures 8A and 8B). If a rater could not make the judgement regarding if a label should be 0 or 1, then the fragment-pair was labeled as NA (See Figure 3D). Such situations typically occurred when one or both fragments in a pair slightly existed outside the OFC, represented cortical folds that were not truly MOS or LOS, or very rarely, sections within a sulcus incorrectly got divided into two fragments due to mis-labeling near the TOS junction. If more than two raters assigned a NA label to a fragment-pair, then we discarded the fragment-pair from the classification analysis. The interrater reliability measured as Cronbach’s alpha for left MOS, left LOS, right MOS, and right LOS, were: 0.9, 0.85, 0.92, and 0.82. In Table 1, we present summaries of total number of sulcal-fragment-pairs that were identified across all subjects and axial-slices.

**Figure 8:**
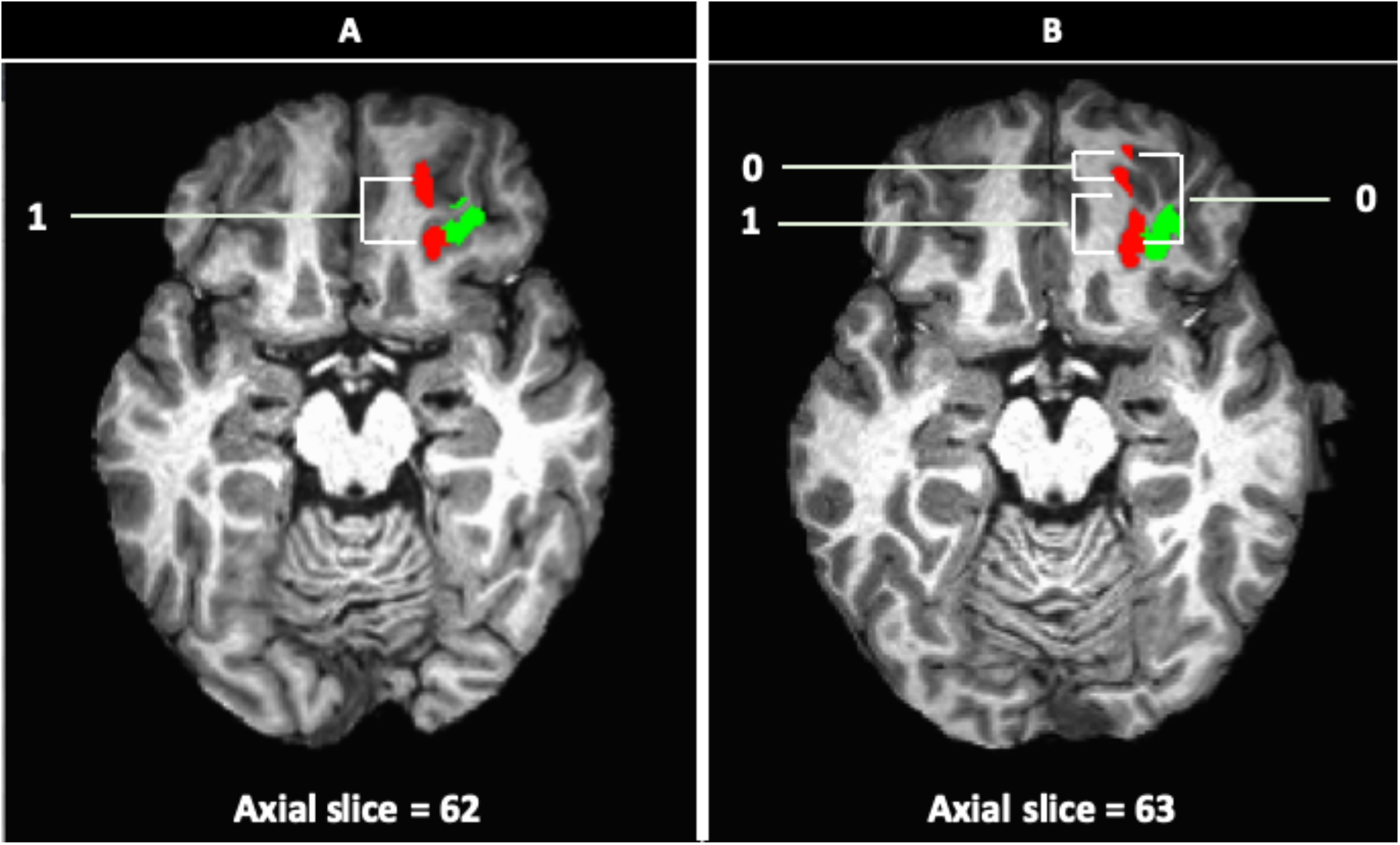
Illustrations of MOS discontinuity within an axial slice. **(A)** The figure illustrates a MOS fragment-pair for which the average rating across the four raters was 1. That is, based on the spatial organization of the MOS fragment-pair, all raters judged the MOS to be discontinuous within the axial slice. **(B)** The figure illustrates 3 pairs of MOS fragments. For two of these fragment-pairs, the average rating across four raters was 0, thus suggesting that the raters could not determine MOS discontinuity within the axial slice by just visualizing those fragment-pairs alone.

**Table 1:** The table summarizes the number of left and right MOS and LOS fragment-pairs with an average rating of: 0, 0.25, 0.33, 0.5, 0.67, 0.75, and 1. All fragment-pairs with average rating below 0.5 were assigned a label 0, and fragment-pairs with average rating above 0.5 were assigned a label 1. The fragments with average rating of 0.5 were discarded from the classification analyses.

|  |  | MOS |  | LOS |  |
| --- | --- | --- | --- | --- | --- |
| Average rating | Final label | Left<br>Count (%) | Right<br>Count (%) | Left<br>Count (%) | Right<br>Count (%) |
| 0.00 | 0 | 443 (58.6%) | 253 (42.5%) | 747 (72.5%) | 321 (50.6%) |
| 0.25 |  | 58 (7.7%) | 52 (8.7%) | 66 (6.4%) | 76 (12.0%) |
| 0.33 |  | 0 (0%) | 0 (0%) | 8 (0.8%) | 2 (0.3%) |
| 0.5 | no consensus | 46 (6.1%) | 45 (7.6%) | 43 (4.1%) | 69 (10.9%) |
| 0.67 |  | 0 (0%) | 2 (0.3%) | 5 (0.5%) | 3 (0.5%) |
| **0.75** | 1 | 84 (11.1%) | 33 (5.5%) | 112 (10.9%) | 99 (15.6%) |
| **1.00** |  | 125 (16.5%) | 210 (35.3%) | 49 (4.8%) | 64 (10.1%) |

##### Classification analysis

We discarded fragment-pairs for which the average value across raters was 0.5 or if at least two raters labeled a fragment-pair as NA, as such cases represented situations where a fragment-pair does not allow consistent judgement of discontinuity across raters. Also, axial slices that contained only one MOS fragment were excluded from this analysis, because to identify discontinuity at least two fragments are needed. We then rounded the averaged labels to 0 or 1 so that a binary classifier can be trained, and the *FV*_*fragment*_ feature vectors along with the binarized labels were saved in a table (See Figure 3E: step-1). We followed the same steps for *FV*_*spine*_ feature vectors, which we saved in a separate table in a similar manner. For a given fragment-pair, the averaged label assigned to its corresponding *FV*_*fragment*_ feature vector, was the same as the label assigned to its *FV*_*spine*_ feature vector.

We conducted two neural network classification experiments using the above tables; the first experiment was conducted using the table of *FV*_*fragment*_ features and the second experiment was conducted using the table of *FV*_*spine*_ features. In each experiment, the columns of the table representing the 8 feature elements were fed to the input layer of the neural network and the last column that contained 0/1 tags were used for guiding the supervised learning. The experiments were run using R’s Neuralnet library (Fritsch, Guenther, Wright, Suling, & Mueller, 2019) (See Figure 3E: step-2). Each neural network experiment consisted of a network with 1 hidden layer, and an output layer with 1 neuron. Based on some initial test runs using a small fraction of our data, the number of hidden neurons were set to 10 to allow reasonably complex decision boundaries to evolve via neural network training. The neurons used a logistic activation function and network weights were adjusted using the rprop+ algorithm. The number of iterations for the learning procedure was set to 200,000 within which the network had to learn to produce an output value less than 0.4 for cases that were tagged as 0, and an output value greater than 0.6 for cases tagged as 1. Each neural network experiment was run in a 5-fold cross-validation style 20 times, and during each run, the classes were balanced using relocating safe-level SMOTE (RSLS) procedure which was available in R’s Smotefamily package (Siriseriwan & Sinapiromsaran, 2016). In all, 200 runs were performed (i.e., 2 experiments x 100 runs); each run yielded the following 6 values:

1. Label-0 prediction accuracy on training data,
2. Label-1 prediction accuracy on training data,
3. Area under the receiver operator characteristic curve (AUC) based on the training data
4. Label-0 prediction accuracy on test data,
5. Label-1 prediction accuracy on test data,
6. AUC based on the test data

For each experiment (i.e., experiment using, *FV*_*fragment*_ or *FV*_*spine*_), the average and the standard deviation of the above 6 values computed across 100 runs were then summarized in a table.

##### Label permutation test

To further test the validity of the features and the associated labels (0/1), for each experiment (i.e., experiment using, *FV*_*fragment*_ or *FV*_*spine*_), we also ran a group-label permutation procedure that consisted of 100 iterations; similar procedures have been implemented previously (Arias, Arratia, & Duarte-López, 2017; Golland & Fischl, 2003; Hsing, Attoor, & Dougherty, 2003; Ojala & Garriga, 2009; Roy et al., 2018). That is, during each iteration, we first balanced the training data using the RSLS procedure, and then randomly 50% cases from class-0 were labeled as 1, and 50% cases from class-1 were labeled as 0 (See Figure 3E: Step-3). Using these permuted labels, we then trained a neural network classifier with the same specifications as the original runs and tested the classifier on the test set whose labels were never permuted. The permutation runs yielded 100 test AUC values which were saved as a vector. To examine the hypothesis that the permuted run’s mean test AUC value across 100 iterations will be significantly lower than the original run’s mean test AUC value across 100 iterations, we performed a 1-sided t-test, which yielded a p-value that we report in a table. In addition to the above, we also report in a table the mean training and test performance of the classifier from the permuted runs.

## 3. Results

The guidelines to characterize MOS or LOS discontinuity were generally informative enough that for most fragment-pairs (~90% to 96%) an overall consensus could be attained (See Table 1). However, it is important to note that consensus was not always 100%, as indicated by some average ratings of 0.25, 0.33, 0.67, or 0.75. We anticipate that this effect might be due to the fact that the spatial organization of MOS and LOS fragment-pairs are quite diverse, and such diversity is not well-captured by illustrative or documentation based guidelines of manual labeling, as a result of which, there is some variability across raters in labeling these cases. The goal of the classification analysis was to thus establish robust decision boundaries to identify MOS or LOS discontinuity while accounting for this spatial diversity in the organization of the fragment-pairs.

For each sulcus, the classification analysis consisted of 2 experiments. In the first experiment, we trained and tested a fixed architecture neural network classifier using *FV*_*fragment*_ features, and in the second experiment, we examined the *FV*_*spine*_ features. The classification analyses results using *FV*_*fragment*_ features are summarized in Table 2, and using *FV*_*spine*_ features are summarized in Table S4. The training accuracies of both classes using *FV*_*fragment*_ features or *FV*_*spine*_ features were typically greater than 99%, and the AUCs were 1. While using the *FV*_*fragment*_ features, the average testing accuracy of label 0 prediction ranged from 95-98% and label 1 prediction ranged from 92-96%, and average AUC was generally greater than 0.98. While using the *FV*_*spine*_ features, the average testing accuracies of label 0 prediction ranged from 94-98% and label 1 prediction ranged from 89-95%, and average AUC was generally greater than 0.98. To further validate the results, we conducted label permutation tests, results of which are summarized in Table 2 and Table S4. As can be seen, our fixed-architecture neural network had considerable trouble in learning the training set when labels were permuted, and hence its performance on the test set, became significantly poor relative to the original runs. Taken together, the above results validated that the feature vectors, *FV*_*fragment*_ and *FV*_*spine*_, did indeed capture the visual features of the sulcal-fragment-pairs that raters relied upon during labeling, and that the framework can learn the mapping between regional cortical folds and predefined human-interpretable shapes by mimicking the overall rating across multiple raters.

**Table 2:** The table illustrates the classification analysis results using the feature vectors, *FV*_*segment*_

|  |  | Training |  |  | Testing |  |  | P-value |
| --- | --- | --- | --- | --- | --- | --- | --- | --- |
|  |  | Accuracy<br>Label-0<br>(mean/se) | Accuracy<br>Label-1<br>(mean/se) | AUC<br>(mean/se) | Accuracy<br>Label-0<br>(mean/se) | Accuracy<br>Label-1<br>(mean/se) | AUC<br>(mean/se) |  |
| Left MOS | Original | 99.7%<br>(0.02) | 99.9%<br>(0.02) | 1.00<br>(0.00) | 97.4%<br>(0.15) | 94.6%<br>(0.35) | 0.99<br>(0.00) | $P < 0.001$ |
|  | Permuted | 45.5%<br>(1.14) | 27.6%<br>(0.88) | 0.65<br>(0.01) | 47.1%<br>(1.10) | 18.9%<br>(1.19) | 0.63<br>(0.00) |  |
| Right MOS | Original | 99.7%<br>(0.04) | 99.6%<br>(0.03) | 1.00<br>(0.00) | 95.6%<br>(0.27) | 95.0%<br>(0.31) | 0.98<br>(0.00) | $P < 0.001$ |
|  | Permuted | 25.4%<br>(0.86) | 71.5%<br>(1.83) | 0.62<br>(0.02) | 28.5%<br>(0.82) | 80.2%<br>(1.13) | 0.68<br>(0.01) |  |
| Left LOS | Original | 99.3%<br>(0.03) | 99.8%<br>(0.02) | 1.00<br>(0.00) | 97.4%<br>(0.11) | 92.7%<br>(0.44) | 0.99<br>(0.00) | $P < 0.001$ |
|  | Permuted | 48.7%<br>(3.95) | 48.8%<br>(3.97) | 0.73<br>(0.02) | 48%<br>(3.84) | 40.6%<br>(3.99) | 0.69<br>(0.02) |  |
| Right LOS | Original | 99.9%<br>(0.01) | 99.8%<br>(0.02) | 1.00<br>(0.00) | 98.0%<br>(0.15) | 95.5%<br>(0.33) | 0.99<br>(0.00) | $P < 0.001$ |
|  | Permuted | 40.4%<br>(1.38) | 26.5%<br>(1.01) | 0.66<br>(0.01) | 40.3%<br>(1.67) | 18.3%<br>(1.27) | 0.63<br>(0.01) |  |

## 4. Discussion

One established way to characterize a regional cortical fold is to visually identify a shape that is embedded within the fold, examine how the embedded shape varies across individuals, and then define manual guidelines to map the structural features of the folding pattern to this distinct set of shape variants. Here, we introduced a novel computational framework such that the manual guidelines can be used to assign labels to example cases of a regional folding pattern based on the shape they represent, and the labels along with feature vectors that capture structural features of the fold, which raters visually rely on, may then be used to train classifiers to learn the mapping. This extension thus eliminates the need for training new human raters and facilitates faster processing of larger datasets. Below, we discuss some important design considerations and the general applicability of the framework.

From a computational viewpoint, the size of the training data and the complexity of the embedded shape within the regional cortical fold are two important factors to consider before implementing the proposed framework. That is, if the human-interpretable shape of interest embedded within the fold is very complex, then to capture its morphology, a feature vector would generally require a greater number of elements relative to shapes that are simpler in nature. Further, training a classifier becomes challenging as the number of variations of the shape increases because, the number represents the distinct classes the classifier has to learn to map the feature vectors to. Thus, for complex shapes, or shapes with multiple variants, or both, it is important to use a dataset with a large number of training cases. With regards to our demonstration problem, we could have designed feature vectors to represent the 4 H-patterns types, which are based on identifying MOS and/or LOS discontinuity within individual axial slices. Instead, we chose to directly address the fundamental issue of identifying MOS and LOS discontinuity, as we anticipated that numerically representing the brokenness of an “H” shape’s vertical arm is relatively simpler than capturing multiple variants of an entire “H” shape. This allowed us to keep the size of our feature vectors reasonably small to 8 elements, each of which represented a visual factor that raters relied upon while characterizing this discontinuity. Also, this decision allowed us to minimize the number of shape variants to 2 (continuous or discontinuous). The total number of feature vectors to train and test the classifiers ranged from 595 to 1307 across the four sulci, which were comparable to the simulation based results of Hua and colleagues (Hua, Xiong, Lowey, Sug, & Dougherty, 2005), which provide guidelines pertaining to optimal feature vector size for a given sample size. In all, by decomposing a complex shape of interest to a set of simpler shapes, we addressed the above design issues. We do however acknowledge that the above steps for constraining the size of feature vectors and the number of classes was specific for the current demonstration problem, and for examining other embedded shapes, alternate approaches may need to be developed to address the above design issues in a problem-specific manner.

The second computational issue pertains to choosing an appropriate representation of regional cortical folds, so that the embedded shape within the fold may be correctly captured. Based on the current literature, computational studies often rely on surface reconstruction procedures to identify voxels that constitute the inner white matter surface, outer gray matter surface (a.k.a the pial surface), and the cortical sheet (Dahnke & Gaser, 2018; Ellis, 2017; Im et al., 2017; Renvall, Witzel, Wald, & Polimeni, 2016; Tran & Fang, 2017). The identified voxels are then processed by mesh generation algorithms (Lederman et al., 2011; Tadel, 2020b, 2020a; Tran & Fang, 2017) to store the cortical voxels as a graph data structure which is then processed by downstream algorithms to identify a connected set of voxels that constitute a variety of sub-structures including sulcal basins (Im et al., 2010; Im, et al., 2013), sulcal-lines (Le Troter, Auzias, & Coulon, 2012), and gyral surface and hinges (Li et al., 2017; Zhang et al., 2018). Alternately, regional cortical folds may also be examined by decomposing a 3D cortical fold into multiple 2D images within axial, coronal, saggital, or 2D oblique plane, and such an approach is ideal for situations where complex regional folding patterns are better examined in a 2D view (Alvarez-Jimenez, Múnera-Garzón, Zuluaga, Velasco, & Romero, 2020; Mangin et al., 2015). For our demonstration, we chose a 2D approach based on our previous determination that available pipelines did not identify all relevant sulci necessary for accurate identification of MOS and LOS discontinuity (Snyder et al., 2019; See Supplement Sections I.d, II.b, and Table S3). Furthermore, analyzing this problem in an axial slice-by-slice manner also allowed us to computationally mimic the existing guidelines pertaining to characterization of MOS and LOS discontinuity. In summary, prior to using the framework, we encourage investigators to conduct preliminary tests to assess in which representation the shapes of interest are visually more apparent and in which the shape can also be encoded as short-size feature vectors.

The computational framework we presented is thus suited for problems similar to our demonstration problem where the underlying spatial organization of different sections of a regional cortical fold need to be examined, and it may be implemented in a 2D or a 3D setting. Further, the elements constituting the feature vectors may utilize conventional measures of regional cortical folds, such as sulcal width and depth (Dahnke, Yotter, & Gaser, 2013; Madan, 2019; Yun, Im, Yang, Yoon, & Lee, 2013), cortical thickness (Dahnke et al., 2013; Tosun, Siddarth, Levitt, & Caplan, 2015), and number of gyral hinges (Li et al., 2009; Li et al., 2017; Zhang et al., 2018), or may be derived using polynomial (Zhang, Guo, Li, Nie, & Liu, 2009) or may be developed using basis function (Rabiei, Richard, Coulon, & Lefèvre, 2019; Yu et al., 2007) based approaches, or other shape-based metrics. We see our framework as a tool for domain experts so that instead of using pictorial illustrations or written guidelines to manually document how embedded patterns within regional cortical folds map to a predefined set of human-interpretable shapes, they can directly use example cases to train classifiers to learn the mapping, which will allow more consistent rating of newer datasets. The framework may also be used for resolving situations where visually similar shapes (e.g., A vs Λ or O vs Θ) appear embedded within a regional fold, but not all variants reliably occur across individuals. We anticipate that such situations are quite common, and impede the discovery of many new embedded patterns in regional cortical folds at various brain regions. In our future work, we hope to test this issue further, and computationally extend this framework by including feature subset selection procedures (Bolón-Canedo & Remeseiro, 2020; Chandrashekar & Sahin, 2014; Hua et al., 2005; Miao & Niu, 2016; Roy, Schaffer, & Laramee, 2015; Way, Sahiner, Hadjiiski, & Chan, 2010; Xue, Zhang, Browne, & Yao, 2016), so that investigators will have added guidance in order to narrow down on the set of physical attributes of a cortical fold that best explain the set of embedded shapes.

## Supporting information

Supplemental_methods_results_and_tables

## 5. Author contributions

A.R., M.P., and V.T. conceptualized and designed the work. A.R. implemented the computational framework. A.R., T.M., D.B., completed quality control and volumetry analyses under the supervision of V.T. A.R., T.M., D.B., W.S. labeled OFC sulcal-fragment-pairs. A.R. and T.M. conducted classification analyses. A.R. and V.T. drafted the article. A.R., T.M., and D.B. worked on the manuscript figures under V.T.’s supervision. A.R., T.M., D.B., W.S., M.P., and V.T. revised the article. All authors approved the final version of the manuscript.

## 6. Acknowledgements

We would like to thank Dr. Brian King, Department of Computer Science, Bucknell University, Lewisburg, PA, 17837 for his valuable comments that helped us improve this work. The data for this work was obtained from the OpenfMRI database (https://openfmri.org/dataset/ds000030/). The work was supported by NIH grant R01 DA44015, which was awarded to Dr. Vanessa Troiani.

## Appendix A. Supplementary information

