## Supplemental_methods_results_and_tables for "A pipeline to characterize local cortical folds by mapping them to human-interpretable shapes"

**Contents**

| | | |
|---|---|---|
| **I** | **Supplementary Methods** | 2 |
| **I.a** | T1 normalization | 2 |
| **I.b** | Estimating whole-brain and local OFC volumes | 3 |
| **I.c** | Developing $FV_{spine}$ features | 4 |
| **I.d** | BrainVisa skeletons versus sulcal-fragment spine | 5 |
| **II.** | **Supplementary Results** | 6 |
| **II.a** | Volumetric analysis | 6 |
| **II.b** | BrainVisa skeletons versus sulcal-fragment spine | 7 |
| **II.c** | Classification results using $FV_{spine}$ features | 7 |
| **III** | **References** | 7 |
| **IV** | **Supplementary Tables** | 9 |
| **S1:** | The results of MRIQC noise analysis | 9 |
| **S2:** | The results of volumetric analysis | 10 |
| **S3:** | The results of BrainVisa skeletons versus sulcal-fragment spine analysis | 12 |

**S4:** The results of classification analysis using $FV_{spine}$ features 13

### I. Supplementary Methods

#### 1.a. T1 normalization

The normalization process involved 4 stages: removing shading artifacts, skull-stripping, affine transformation to MNI-space, and non-linear warping to an AFNI template in MNI space. Below, we discuss the processing involved in each of these stages in detail.

***Removing shading artifacts:*** Each subject-specific T1 weighted image was processed using AFNI's 3dUnifize function to remove shading artifacts; the procedure is recommended as a preprocessing step prior to skull-stripping (*AFNI program: 3dUnifize*, 2020; Cox, 1996). For this procedure, the input parameters were set to default values. For each subject, the function produced an image for which the white matter intensity was spatially more uniform than the raw image.

***Skull-stripping:*** In the next step, the 3dunifized image of each subject was processed using AFNI's 3dSkullStrip function to exclude the skull from the image. Although, 3dskullStrip function may be run in a single step to obtain a 3D-brain image with skull excluded, we found that this way of running the procedure introduced some unwanted spatial smoothing. To circumvent this issue, for each subject, we chose to run the procedure in two stages. First, using 3dSkullstrip function, a binary mask was generated representing the in-brain voxels (i.e., skull excluded). The mask was then applied to the 3dunifized T1 image to obtain the final skull strip image. The above way of running the procedure thus guaranteed that the final image for each subject was derived directly from the input image and that no voxels were smoothed. For this step the number of iterations was set to 400 and the parameter, *LD*, that defines the surface density, was set to 40. The above parameters were set empirically based on multiple initial test runs to examine what setting resulted in the most accurate identification of the skull. For each subject, the output of this stage was a skull-stripped NIFTI image in native space.

***Affine transformation:*** In the third stage, each skull-stripped image was affine transformed to MNI space using AFNI's 3dAllineate function to match AFNI's MNI152_2009 skull-stripped template (*SSwarper base volumes*, 2020). The general idea behind affine transformation is to match a subject's brain to the MNI template brain at a whole-brain-level; the step does not guarantee alignment of finer anatomical structures. For this step, we chose the option of two-pass alignment with absolute local Pearson's correlation as the cost function. The explicit smoothing parameter "fineblur" was set to 0 mm, and the parameter "cmass" was set to determine an initial shift to align the images based on a center-of-mass calculation. For each subject, the output of this stage was a NIFTI image in MNI-space of dimension, 193x229x193 mm.

***Non-linear warping:*** In the final stage, the objective was to match each subject's affine transformed image to AFNI's MNI152_2009 skull-stripped template (*SSwarper base volumes*, 2020) using non-linear warping so that the local anatomical structures are better matched between the images. A prior simulation study using 101 T1 images from MindBoggle dataset compared non-linear warping procedures such as, ANT, DARTEL, FNIRT, and 3dqwarp, in terms of their ability to preserve known spatial structures in space (Cox & Glen, 2013). In this study, the 3dqwarp transformed images showed 80-90% overlap of OFC-labeled voxels across subjects, which was higher than the overlap values obtained using the other procedures. Based on these findings, we chose the 3dqwarp procedure to perform non-linear warping. For this procedure, we set the explicit blurring related parameters to 0 mm. For the cost function, we used the default option of clipped Pearson correlation. Based on some initial test runs, we set the minimum patch size for warp searching to 19 mm. For each subject, the output of this stage was a NIFTI image in MNI-space of dimension, 193x229x193 mm. We visually inspected each normalized image for non-linear distortions (e.g., over smoothing, stretching) within OFC. The distorted cases were discarded from further analyses.

**I.b. Estimating whole-brain and local OFC volumes**

DARTEL utilized each subject's T1 image in native space and performed the following 4 steps. In step-1, it extracted gray matter, white matter, and cerebral spinal fluid (CSF) and saved them in NIFTI format in native space. In step-2, it normalized the gray matter tissue probability map to MNI space. Then in step-3, it modulated the normalized gray matter image to preserve tissue volume. Finally, in step-4, it smoothed the normalized-modulated gray matter image using a Gaussian kernel with full-width at half maximum value of 4 mm to produce smwc* file. Thus for each subject, the smwc* file existed in MNI-space, and were then saved in NIFTI format. Using the files generated from step-1 and step-4, we then computed global and local volumes as follows:

*A. Estimation of whole-brain volumes:* The whole-brain gray-matter, white-matter, and CSF volumes in milliliters (ml) were estimated using the gray matter segmented image produced in step-1. The value of each voxel in this image represented the fraction of the voxel's volume that contained a specific tissue type (gray, white, or CSF), and it ranged from 0 to 1. In these images, voxels with value less than 0.2 were set to 0 as they represented noisy voxels. Our choice for this threshold was based on the study by Callaert, Ribbens, Maes, Swinnen, and Wenderoth (Callaert, Ribbens, Maes, Swinnen, & Wenderoth, 2014). The total gray-matter volume was then defined as the sum across all voxels in the thresholded gray-matter image multiplied by the volume of a voxel in milliliters (ml). The whole-brain white-matter and CSF volumes were estimated using the corresponding threshold files by following the same steps. Our whole-brain volumetry results were comparable to the volumes of healthy subjects as reported in the published literature (Brown et al., 2011; Callaert, Ribbens, Maes, Swinnen, & Wenderoth, 2014; Kim, Kim, & Jeong, 2017).

*B. Estimation of OFC gray-matter volume:* For each subject, as the modulation step preserves the gray matter volume both globally and locally, the sum of all voxels in smwc* file multiplied by a voxel's volume in the smwc* file is approximately equal to the overall gray matter volume estimated in the native space (See paragraph above). In other words, the smwc* file provides us an alternate way to compute a subject's whole-brain gray matter volume. An added advantage of this approach is that by using existing atlases, which are in MNI-space, we can also compute local brain volumes (e.g., OFC volume). One challenge however is to establish an optimal gray matter probability threshold for the smwc* file in a subject-specific manner so that the whole-brain gray matter volume computed using the smwc file may match very closely (e.g., $2^{nd}$ decimal point) to the whole-brain gray matter volume computed in the native space. To estimate such an optimal gray matter probability threshold, we implemented a 1-D adaptive procedure in MATLAB as follows. For a given threshold value, the procedure computes the squared error between overall gray matter volume in the native space (native space image threshold is fixed at 0.2) and the gray matter volume derived using the smwc* file by discarding voxels with values lower than the threshold. The procedure then systematically varies the threshold, and the threshold at which the error minimizes is chosen as the optimal threshold.

The procedure was run in a subject-specific manner to derive an individual's smwc* probability threshold. For each subject, we then overlaid the AAL atlas (Rolls, Huang, Lin, Feng, & Joliot, 2020; Tzourio-Mazoyer et al., 2002) on their respective thresholded smwc* file. Using a custom-made MATLAB routine, the voxels that constituted the left and right OFC were identified. The left and right OFC volumes were then defined as the total sum across all voxel values in the left and right OFC regions of the thresholded smwc* file, respectively, multiplied by the volume of a voxel in the smwc file. We ran the above steps for all subjects, and saved the global gray-matter, white-matter, and CSF volumes, and local left and right hemisphere OFC volumes in milliliters (ml) in a table.

#### I.c. Developing $FV_{spine}$ features

For spine detection we used a betweenness centrality type approach which is similar to a procedure that Le Troter, Auzias, and Coulon (Le Troter, Auzias, & Coulon, 2012) used for generating a line representation of a sulcus using a 3D mesh graph of a cortical fold. In contrast to Le Troter, Auzias, and Coulon's approach, our custom-made procedure was designed specifically to function in a 2D image space instead of a 3D mesh space. In our implementation, we used a moving window type approach that allowed us to effectively capture the shape of a fragment of a cortical fold in a 2D plane which is reasonably orthogonal to the fold. Below, we discuss the details of the procedure for a MOS or a LOS fragment.

Given a 2D MOS or a LOS fragment (See section 2.2.3), the procedure first establishes a directional lattice graph such that each voxel (voxels = graph nodes) constituting the fragment is connected to 4 neighboring voxels in space. The connection weight from voxel-i to voxel-j is defined as the image intensity value of voxel-j, and the connection weight from voxel-j to voxel-i is defined as the image intensity value of voxel-i. We then define a moving rectangular window of size 10 voxels in posterior-to-anterior direction; we set the window size based on some initial experimental runs. The procedure then aligns the posterior part of the window with the posterior-most part of the fragment, and then extracts a sub-graph using voxels that exists within the window. For each node of this sub-graph, betweenness centrality is computed; the betweenness centrality of a node in a graph represents the number of shortest paths between all pairs of nodes in the graph that pass through the node. Since the weights of edges going towards low image intensity voxels are very low, a considerable number of shortest paths in the sub-graph passes through such voxels. The procedure then repeats the above steps by moving the window by 5 voxels in the anterior direction each time. The average betweenness centrality value of each voxel across the moving window is then computed and all voxels with average betweenness centrality value greater than 0.3 are then chosen to represent the fragment's spine. The parameters, window step-size (i.e., 5 voxels) and betweenness centrality threshold of 0.3 are based on some initial test runs where we examine if the spines generated by the above procedure captures the shape of the test fragments reasonably well. An example output of the procedure is shown in Figure 7. That is, in Figure 7A, we show an axial slice of a T1 image. Then in Figure 7B, we show the MOS, TOS, LOS, and IOS fragments, and in Figure 7C, we illustrate the spine of MOS, TOS, and LOS fragments. As can be seen, the spine summarizes the general structure of a fragment of cortical fold in a 2D plane much like how a non-linear regression line summarizes a scatter plot. Thus, a fragment is analogous to a canyon, and a spine is a river that flows through the canyon.

**I.d. BrainVisa skeleton versus sulcal-fragment spine analysis**

We ran BrainVISA's Morphologist-2015 pipeline on the AFNI normalized T1 images of our subjects to derive their sulcal skeletons; no additional normalization was done during this processing and all numeric parameters were set to their default values. Based on an initial visual examination, we found that for several MOS, TOS, and LOS fragments, no skeletons were identified by the BrainVISA pipeline. This effect might have been due to the fact that line representations in 3D mesh space are based on identifying a path across mesh-nodes that represent image minimas across OFC axial slices, and might have therefore, under-represented image minimas within axial slices. Using a custom-made R-based procedure, for left MOS, right MOS, left LOS, and right LOS, we additionally compared spines and skeletons using the following five metrics: (1) the number of sulcal-fragments that contained these structures (spines or skeletons), (2) the mean sizes of spines and skeletons measured in voxels, (3) the mean overlap between the voxels of spines and skeletons within a given fragment measured as dice coefficient, (4) the minimum, mean, and maximum distances between the voxels of spines and skeletons within a sulcal-fragment, and (5) the mean normalized cumulative intensity of spines and skeletons measured as the ratio of the sum of image intensities of all voxels constituting spine or skeletons divided by the 95$^{th}$ percentile image intensity of the axial slice. The rationale for normalizing the image intensity was that the range of numeric values of voxels after normalization varied across images. Also, we chose not to use the maximum for normalization because we anticipated that such an isolated high value within an axial slice might be an outlier value, and therefore, we used the image intensity value at 95$^{th}$ percentile. The measures 3 and 4 were computed using sulcal-fragments that contained both spines and skeletons. We present the results of BrainVISA skeleton versus sulcul-fragment spine analysis in Table S4, and discuss the results in Supplemental Section II.b. In all, the results suggested that both spines and BrainVISA skeletons represented spatially close low intensity voxels.

### II. Supplementary Results

#### II.a Volumetric analysis

To identify cases with abnormal local OFC volume, we performed volumetric analyses using SPM12's DARTEL toolbox. The subject-specific volumetric values are reported in Supplementary Table S2. We first compared the global brain volumes of the subjects that passed the image quality test to the global brain volumes of healthy control subjects reported in recent literature. For our cases that passed MRIQC analysis, the mean whole-brain gray-matter volume was 687.8 ml (sd = 66.4), white-matter volume was 447.3 ml (sd = 48.9), and CSF volume was 255.2 ml (sd = 68.1). These values are comparable to the whole brain volumes of healthy subjects reported in the literature (Brown et al., 2011; Callaert et al., 2014; Kim, Kim, & Jeong, 2017). For example, in a study involving healthy subjects (Callaert et al., 2014), the mean gray-matter, the mean white-matter, and the mean CSF volumes were reported as, 688 ml (sd = 57.0), 489 ml (sd = 42.8), and 255 ml (sd = 18.1), respectively.

In the next step, we examined these measures at a group level to identify outlier cases using box plot analyses. As no outliers were detected at a global whole-brain level, we conducted 2 local OFC level analyses, one for each hemisphere. The mean local OFC gray matter volume for left hemisphere was 15.2 ml (sd = 1.9) and for the right hemisphere was 15.3 ml (sd = 2.0). The local analyses found no outliers beyond 1.5 times the interquartile range above and below the upper and lower quartile, respectively. The above findings thus suggested that the global brain volumes of the subjects we used for the classification analyses were consistent with the brain volumes of healthy subjects as reported in the published literature, and that there were no outlier local OFC volume cases.

**II.b. BrainVisa skeleton versus sulcal-fragment spine analysis**

In Table S4, we summarize the results of BrainVisa skeleton versus sulcal-fragment spine analysis. As shown in the table, the total number of sulcal-fragments identified for the four sulci varied between 700 to 1200. For LOS, the number of 2D sulcal-fragments was generally greater than MOS. Our custom-made spine detection procedure (Section I.c) was able to generate spines for more than 99% of these fragments. In contrast, BrainVisa based skeletons estimated using default BrainVisa parameter settings was able to identify skeletons for approximately 55% of the left MOS, left and right LOS fragments, and about 65% of right MOS fragments. The sizes of the sulcal fragments varied from 28 to 54 voxels, sizes of the spines varied from 11 to 22 voxels, and the sizes of the skeletons varied from 5 to 13 voxels. Although, the overlap between the spines and the skeletons in terms of Sorenson's score was low and was typically around 0.32, the mean distance between their voxels was generally less than 4 mm, and their mean normalized cumulative intensities were reasonably similar, thus suggesting that they represented spatially close low intensity voxels.

**II.c Classification results using $FV_{spine}$ features**

Using $FV_{spine}$ feature table, we ran 4 classification analyses: one for left MOS, one for right MOS, one for left LOS, and one for right LOS.. Each analysis was run in a 5-fold crossvalidation format 20 times. In all, for each analysis, there were 100 runs. The results of the classification analysis using $FV_{spine}$ features are summarized in Table S3. As shown in the table, the average training accuracy of label 0 and label 1 prediction across 100 runs was greater than 99% and the training AUCs for all the four runs was 1. For these runs, the average testing accuracies of label 0 prediction ranged from 94-96% and label 1 prediction ranged from 83-94%, and average test AUCs were generally greater than 0.96.

### IV. Supplemental Tables

**Table S1:** The table summarizes the results of MRIQC noise analysis

| Sr. No. | Subjects | CJV | Comments |
|---|---|---|---|
| **1** | 10159 | **0.51** | The Image quality was poor in both native space and the MNI space |
| **2** | 10217 | 0.39 | |
| **3** | 10290 | 0.43 | |
| **4** | 10304 | 0.45 | |
| **5** | 10321 | 0.43 | |
| **6** | 10325 | 0.46 | |
| **7** | 10329 | 0.42 | |
| **8** | 10339 | 0.44 | |
| **9** | 10356 | 0.42 | |
| **10** | 10365 | 0.42 | Noise in OFC region in MNI space |
| **11** | 10376 | **0.51** | The Image quality was poor in both native space and the MNI space |
| **12** | 10388 | 0.44 | |
| **13** | 10429 | 0.46 | |
| **14** | 10440 | 0.44 | |
| **15** | 10487 | 0.45 | |
| **16** | 10492 | 0.4 | |
| **17** | 10517 | 0.39 | |
| **18** | 10557 | 0.4 | |
| **19** | 10570 | 0.41 | |
| **20** | 10575 | 0.44 | The OFC was noisy after normalization |
| **21** | 10624 | 0.39 | |
| **22** | 10631 | 0.43 | |
| **23** | 10638 | 0.41 | |
| **24** | 10668 | 0.43 | |
| **25** | 10672 | 0.41 | |
| **26** | 10680 | 0.39 | |
| **27** | 10696 | 0.4 | |
| **28** | 10708 | 0.42 | |
| **29** | 10724 | 0.34 | |
| **30** | 10779 | 0.43 | |
| **31** | 10785 | 0.38 | |
| **32** | 10877 | 0.46 | |
| **33** | 10882 | 0.46 | |

| | | | |
|---|---|---|---|
| **34** | 10912 | 0.38 | |
| **35** | 10934 | 0.38 | |
| **36** | 10940 | 0.39 | |
| **37** | 10968 | 0.42 | |
| **38** | 10975 | 0.45 | |
| **39** | 10977 | 0.47 | |
| **40** | 11019 | 0.44 | |
| **41** | 11050 | 0.48 | |
| **42** | 11059 | 0.43 | |
| **43** | 11066 | 0.44 | |
| **44** | 11077 | 0.39 | |
| **45** | 11082 | 0.39 | |
| **46** | 11088 | 0.42 | |
| **47** | 11097 | 0.46 | |
| **48** | 11104 | 0.42 | |
| **49** | 11105 | 0.43 | |
| **50** | 11112 | **0.50** | The Image quality was poor in the native space |
| **51** | 11131 | 0.45 | |
| **52** | 11143 | 0.38 | |

**Table S2:** The table summarizes the results of volumetric analysis. The columns gray, white, and CSF, represent the gray matter, white matter, and CSF volumes in milliliter (ml) in native space. The column gray-MNI represents the gray matter volume computed using the smwc* files in MNI space. The subject-specific 1-dimensional adaptive procedure allowed us to tune the smwc* probability thresholds such that the gray matter volume computed in the native space matched to the gray matter volume computed in MNI space using the smwc* file.

| Sr. no. | Subject | Gray (ml) | White (ml) | CSF (ml) | Gray-MNI (ml) | Orbital-left (ml) | Orbital-right (ml) |
|---|---|---|---|---|---|---|---|
| **1** | 10159 | 655.7 | 418.8 | 201.8 | 655.7 | 14.4 | 14.0 |
| **2** | 10217 | 648.5 | 462.7 | 188.1 | 648.5 | 15.3 | 15.6 |
| **3** | 10290 | 593.0 | 417.6 | 348.4 | 593.0 | 12.3 | 12.2 |
| **4** | 10304 | 786.0 | 511.2 | 269.4 | 786.0 | 17.9 | 17.6 |
| **5** | 10321 | 673.4 | 527.4 | 254.9 | 673.4 | 14.7 | 14.8 |
| **6** | 10325 | 640.6 | 378.9 | 196.7 | 640.6 | 13.3 | 13.6 |
| **7** | 10329 | 733.0 | 501.9 | 214.5 | 733.0 | 16.0 | 15.9 |
| **8** | 10339 | 589.2 | 388.9 | 268.2 | 589.2 | 12.5 | 12.3 |
| **9** | 10356 | 679.5 | 440.0 | 291.7 | 679.5 | 16.1 | 16.7 |

| | | | | | | | |
|---|---|---|---|---|---|---|---|
| **10** | 10365 | 747.3 | 473.1 | 183.4 | 747.3 | 16.6 | 17.3 |
| **11** | 10376 | 661 | 527.4 | 299.5 | 661.0 | 15.6 | 14.9 |
| **12** | 10388 | 577.4 | 349.4 | 278.7 | 577.4 | 12.0 | 11.9 |
| **13** | 10429 | 636.3 | 399.8 | 169.1 | 636.3 | 12.8 | 13.4 |
| **14** | 10440 | 696.7 | 484.2 | 284.7 | 696.7 | 15.1 | 16.0 |
| **15** | 10487 | 668.8 | 456.4 | 321.1 | 668.8 | 15.1 | 14.9 |
| **16** | 10492 | 633.9 | 373.1 | 429.6 | 633.9 | 14.0 | 13.9 |
| **17** | 10517 | 815.7 | 478.4 | 192.7 | 815.7 | 19.6 | 20.1 |
| **18** | 10557 | 743 | 442.8 | 364.8 | 743.0 | 17.3 | 17.2 |
| **19** | 10570 | 742.8 | 511.6 | 334.1 | 742.8 | 17.0 | 17.5 |
| **20** | 10575 | 768.5 | 460.1 | 251.4 | 768.5 | 17.0 | 17.7 |
| **21** | 10624 | 814.5 | 510.9 | 245.1 | 814.5 | 18.6 | 18.5 |
| **22** | 10631 | 649.2 | 432.2 | 177.5 | 649.2 | 14.2 | 14.3 |
| **23** | 10638 | 607.2 | 448.5 | 283.4 | 607.2 | 12.9 | 12.9 |
| **24** | 10668 | 709.2 | 428.2 | 246.0 | 709.2 | 16.2 | 15.9 |
| **25** | 10672 | 653.3 | 447.3 | 231.6 | 653.3 | 14.1 | 14.5 |
| **26** | 10680 | 746.8 | 483.3 | 179.6 | 746.8 | 18.0 | 18.1 |
| **27** | 10696 | 772.2 | 480.2 | 269.7 | 772.2 | 17.6 | 18.1 |
| **28** | 10708 | 698.8 | 468.2 | 211.2 | 698.8 | 15.4 | 14.5 |
| **29** | 10724 | 772.5 | 396.9 | 177.4 | 772.5 | 17.7 | 18.0 |
| **30** | 10779 | 620.8 | 360.3 | 161.3 | 620.8 | 13.5 | 13.6 |
| **31** | 10785 | 676.3 | 390.3 | 162.2 | 676.3 | 14.8 | 14.7 |
| **32** | 10877 | 615.1 | 425.1 | 315.2 | 615.1 | 12.5 | 12.4 |
| **33** | 10882 | 688.4 | 471.7 | 226.4 | 688.4 | 14.3 | 14.1 |
| **34** | 10912 | 645.2 | 415.1 | 233.5 | 645.2 | 14.9 | 15.0 |
| **35** | 10934 | 654.9 | 435.9 | 253.7 | 654.9 | 14.3 | 13.7 |
| **36** | 10940 | 744.2 | 518.1 | 402.6 | 744.2 | 17.1 | 17.2 |
| **37** | 10968 | 757.2 | 517.6 | 195.1 | 757.2 | 15.8 | 15.8 |
| **38** | 10975 | 710.2 | 524.7 | 365.2 | 710.2 | 15.4 | 16.1 |
| **39** | 10977 | 594.6 | 411.4 | 290.5 | 594.6 | 12.5 | 12.4 |
| **40** | 11019 | 659.6 | 383.3 | 261.8 | 659.6 | 14.9 | 14.5 |
| **41** | 11050 | 701.1 | 470.7 | 266.9 | 701.1 | 14.8 | 15.4 |
| **42** | 11059 | 721.3 | 406.5 | 233.1 | 721.3 | 16.1 | 16.5 |
| **43** | 11066 | 682.3 | 494.1 | 338.7 | 682.3 | 14.5 | 14.6 |
| **44** | 11077 | 696.4 | 439.2 | 199 | 696.4 | 14.7 | 15.0 |
| **45** | 11082 | 641.7 | 461.1 | 252.3 | 641.7 | 14.6 | 14.6 |
| **46** | 11088 | 702.1 | 418.2 | 252.1 | 702.1 | 15.4 | 15.4 |
| **47** | 11097 | 636.6 | 446.3 | 328.4 | 636.6 | 13.4 | 13.9 |
| **48** | 11104 | 813.4 | 461.7 | 226.7 | 813.4 | 18.3 | 18.6 |

| | | | | | | | |
|---|---|---|---|---|---|---|---|
| **49** | 11105 | 649.0 | 438.5 | 188.9 | 649.0 | 14.4 | 14.8 |
| **50** | 11112 | 696.7 | 417.1 | 234.1 | 696.7 | 15.3 | 15.6 |
| **51** | 11131 | 533.8 | 345.4 | 127.8 | 533.8 | 11.5 | 11.5 |
| **52** | 11143 | 759.0 | 530.0 | 362.0 | 759.0 | 16.9 | 17.0 |

**Table S3:** The results of BrainVISA skeleton versus sulcal-fragment spine analysis. The skeletons were generated using BrainVISA Morphologist 2015 pipeline.

| **Measures** | | **Left MOS** | **Right MOS** | **Left LOS** | **Right LOS** |
|---|---|---|---|---|---|
| **Total (%)** | Sulcal fragment | 782 | 709 | 1158 | 854 |
| | Spine | 780 (99.7%) | 708 (99.9%) | 1153 (99.6%) | 852 (99.8%) |
| | Skeleton | 425 (54.3%) | 462 (65.2%) | 635 (54.8%) | 458 (53.6%) |
| **Size** (Voxels) | Sulcal fragment | 28.74 (0.89) | 47.71 (1.28) | 63.89 (2.10) | 53.20 (1.78) |
| | Spine | 11.59 (0.38) | 18.20 (0.49) | 21.63 (0.61) | 19.08 (0.56) |
| | Skeleton | 5.13 (0.19) | 5.43 (0.18) | 12.53 (0.46) | 8.63 (0.29) |
| **Overlap** (Sorenson score) | Spine vs skeleton | 0.36 (0.01) | 0.29 (0.01) | 0.32 (0.01) | 0.31 (0.01) |
| **Spine-to-skeleton distance** (mm) | Minimum | 0.05 (0.01) | 0.06 (0.01) | 0.06 (0.01) | 0.05 (0.01) |
| | Mean | 3.16 (0.10) | 3.94 (0.10) | 5.46 (0.12) | 4.69 (0.10) |
| | Maximum | 7.76 (0.27) | 9.78 (0.28) | 14.09 (0.36) | 11.89 (0.30) |
| **Normalized cumulative intensity** | Sulcal fragment | 15.16 (0.46) | 24.45 (0.63) | 31.55 (1.02) | 26.40 (0.87) |

| | | | | | |
|---|---|---|---|---|---|
| | Spine | 5.59 (0.17) | 8.44 (0.22) | 9.32 (0.25) | 8.34 (0.23) |
| | Skeleton | 2.33 (0.08) | 2.29 (0.08) | 4.67 (0.18) | 3.12 (0.11) |

**Table S4:** The table illustrates the classification analysis results using the feature vectors, $FV_{spine}$

| | | Training | | | Testing | | | |
|---|---|---|---|---|---|---|---|---|
| | | **Accuracy Label-0** (mean/se) | **Accuracy Label-1** (mean/se) | **AUC** (mean/se) | **Accuracy Label-0** (mean/se) | **Accuracy Label-1** (mean/se) | **AUC** (mean/se) | **P-value** |
| **Left MOS** | Original | 99.5% (0.03) | 99.6 (0.02) | 1.00 (0.00 ) | 95.7% (0.18) | 91.1% (0.42) | 0.98 (0.00) | *P < 0.001* |
| | Permuted | 48.5% (1.22) | 31.9% (1.03) | 0.68 (0.01) | 46.4% (1.07) | 22.7 (1.17) | 0.62 (0.01) | |
| **Right MOS** | Original | 99.3% (0.05) | 99.6% (0.02) | 1.00 (0.00) | 94.0% (0.31) | 93.8% (0.32) | 0.98 (0.00) | *P < 0.001* |
| | Permuted | 28.7% (0.97) | 69.7% (1.81) | 0.65 (0.02) | 32.0% (1.04) | 77.3% (1.17) | 0.72 (0.01) | |
| **Left LOS** | Original | 99.6% (0.02) | 99.8% (0.01) | 1.00 (0.00) | 97.0% (0.12) | 89.1% (0.52) | 0.98 0.00) | *P < 0.001* |
| | Permuted | 37.7% (2.81) | 36.8% (2.86) | 0.69 (0.02) | 35.4% (2.78) | 24.9% (2.93) | 0.58 (0.02) | |
| **Right LOS** | Original | 99.8% (0.02) | 99.9% (0.01) | 1.00 (0.00) | 97.8% (0.13) | 94.8% (0.34) | 0.99 (0.00) | *P < 0.001* |
| | Permuted | 47.7% (1.47) | 33.4% (1.09) | 0.68 (0.01) | 46.7% (1.49) | 24.5 (1.25) | 0.62 (0.01) | |

.
